# Fast scanned widefield scheme provides tunable and uniform illumination for optimized SMLM on large fields of view

**DOI:** 10.1101/2020.05.08.083774

**Authors:** Adrien Mau, Karoline Friedl, Christophe Leterrier, Nicolas Bourg, Sandrine Lévêque-Fort

## Abstract

Quantitative analyses in classical fluorescence microscopy and Single Molecule Localization Microscopy (SMLM) require uniform illumination over the field of view; ideally coupled with optical sectioning techniques such as Total Internal Reflection Fluorescence (TIRF) to remove out of focus background. In SMLM, high irradiances (several kW/cm^²^) are crucial to drive the densely labeled sample into the single molecule regime, and conventional gaussian-shaped lasers will typically restrain the usable field of view to around 40 µm x 40 µm. Here we present Adaptable Scanning for Tunable Excitation Regions (ASTER), a novel and versatile illumination technique that generates uniform illumination over adaptable fields of view and is compatible with illumination schemes from epifluorescence to speckle-free TIRF. For SMLM, ASTER delivers homogeneous blinking kinetics at reasonable laser power, providing constant precision and higher throughput over fields of view 25 times larger than typical. This allows improved clustering analysis and uniform size measurements on sub-100 nm objects, as we demonstrate by imaging nanorulers, microtubules and clathrin-coated pits in COS cells, as well as periodic β2-spectrin along the axons of neurons. ASTER’s sharp, quantitative TIRF and SMLM images up to 200 µm x 200 µm in size pave the way for high-throughput quantification of cellular structures and processes.

## Introduction

In advanced widefield fluorescence microscopy, lasers are a common excitation source: they provide excitation at precise wavelengths and possess high spatial coherence, both properties that are crucial to obtain quantifiable images. Typically, the laser is focused at the Back Focal Plane (BFP) of the objective to produce a large collimated beam illuminating the whole sample. While widefield fluorescence microscopy is a fast imaging method, resulting images are usually contaminated by blur from below and above the plane of focus, clouding the fluorescence signal.

Optical sectioning improves the signal by spatially limiting the illumination around the focal plane: Highly Inclined and Laminated Optical Sheet (HiLo) excitation translates the laser beam in the BFP to an oblique illumination of the sample^1^. Placing the beam at the position corresponding to illumination at the critical angle results in a “grazing incidence”, ∼1-µm thick illumination sheet above the coverslip^2,3^. Inclining the beam further^4^ results in Total Internal Reflection Fluorescence (TIRF^5^), restraining the illumination to an exponentially decreasing intensity over a few tens of nanometers above the coverslip surface. These remarkable sectioning capabilities can be performed on one single setup^6–8^ and allow to study membrane and adhesion processes with minimal background. In practice, however, TIRF suffers from heterogeneous illumination caused by the interference patterns arising from the high spatial coherence of lasers and scattering. Rapidly spinning the beam around the BFP can alleviate these fringes by averaging beam orientations over a single camera frame^9^, a method since applied with several variants and refinements^10–12^.

The methods above result in Gaussian-shaped illumination profiles over the sample. This is sufficient for the typical Field of View (FOV) acquired by EMCCD cameras but is more problematic over larger FOVs acquired by newer, highly sensitive sCMOS cameras^13^. The non-uniformity of Gaussian-shaped illumination lowers exploitable FOV sizes and thereby decreases the imaging throughput, a significant caveat for quantitative analysis of images obtained by TIRF.

The need for uniform excitation is even more pressing in Single Molecule Localization Microscopy (SMLM) such as (f)PALM^14,15^ or PAINT^16^ where the localization precision strongly varies with the emitted photons. It’s even more dramatic in (d)STORM^17–19^ f where the single molecule regime (≪1 emitting molecule/µm^3^) relies on driving most fluorophores in a dark state, provided through a high irradiance (kW/cm^2^). As the transition to the dark state is highly dependent on the local excitation intensity, non-uniformities of illumination result in a strongly heterogeneous blinking behavior and loss of image quality across the FOV. For all SMLM methods, no matter the origin of the single molecule emission, non-uniform precision precludes proper analysis of SMLM images over large fields of view.

Thus, several recent studies have aimed at obtaining a uniform excitation over a large FOV. For example, wave-guides^20–22^, provide excellent fixed TIRF on extremely large fields, but cannot be restricted to the actual FOV acquired by the camera, illuminating and bleaching the whole sample at once. Classical solutions revolve around beam-reshapers^23–26^ and multimode fibers^27–29^ but are also extremely restrained in field adaptability. Spatial light modulators (piSMLM^30^) may adapt shape and size of the FOV but suffers high power loss and is rather expensive and complex. All of these classical methods illuminate the whole field at once, so they may be ill-adapted to TIRF (Supplementary Note 1), need speckle reducers and provide larger FOVs under the premise of using higher laser power. However, focusing a high-power laser beam at the edge of the BFP may damage the lens at the back of the objective.

To circumvent the compromise between laser power requirements, optical sectioning performances and field uniformity, we developed ASTER (Adaptable Scanning for Tunable Excitation Region). ASTER is a hybrid scanning and widefield excitation scheme that can perform epifluorescence, oblique or TIRF illumination, while providing illumination uniformity at variable FOV sizes adapted to the camera or sample. Being a general widefield illumination scheme, ASTER can benefit to both classical widefield fluorescence microscopy and SMLM.

### Flat-top epifluorescence/TIRF excitation

ASTER is a hybrid scanning and widefield excitation scheme. Any classical wide field setup can be converted into ASTER configuration by smoothly integrating alternative optical conjugation along with a scanning device such as galvanometers. In our implementation (Fig. 1a) the initial gaussian beam, which provides a limited and non-uniform excitation is focalized between two galvanometer scanning mirrors placed in a plane conjugated to the BFP of the objective so that an angle shift applied to the mirrors will induce a similar angle shift in the objective BFP and a position shift of the beam at sample plane. This configuration allows for large XY area scans of a collimated beam. Fast scanning of the Gaussian beam position in defined patterns such as raster scan or an Archimedes spiral then generates an overall homogeneous illumination (Supplementary Fig. 1) over the FOV when averaged over the camera frame exposure time. Interestingly, field size can be increased or diminished in milliseconds without physical intervention by adapting the galvanometer input amplitude. Notably, as the polar angle of the beam varies at the BFP while its position is maintained, this flat-top excitation scheme is compatible with inclined illumination such as oblique or TIRF. To this end, a conventional motorized translation stage serves as switch from epifluorescence to oblique and TIRF excitation.

**Fig. 1:**
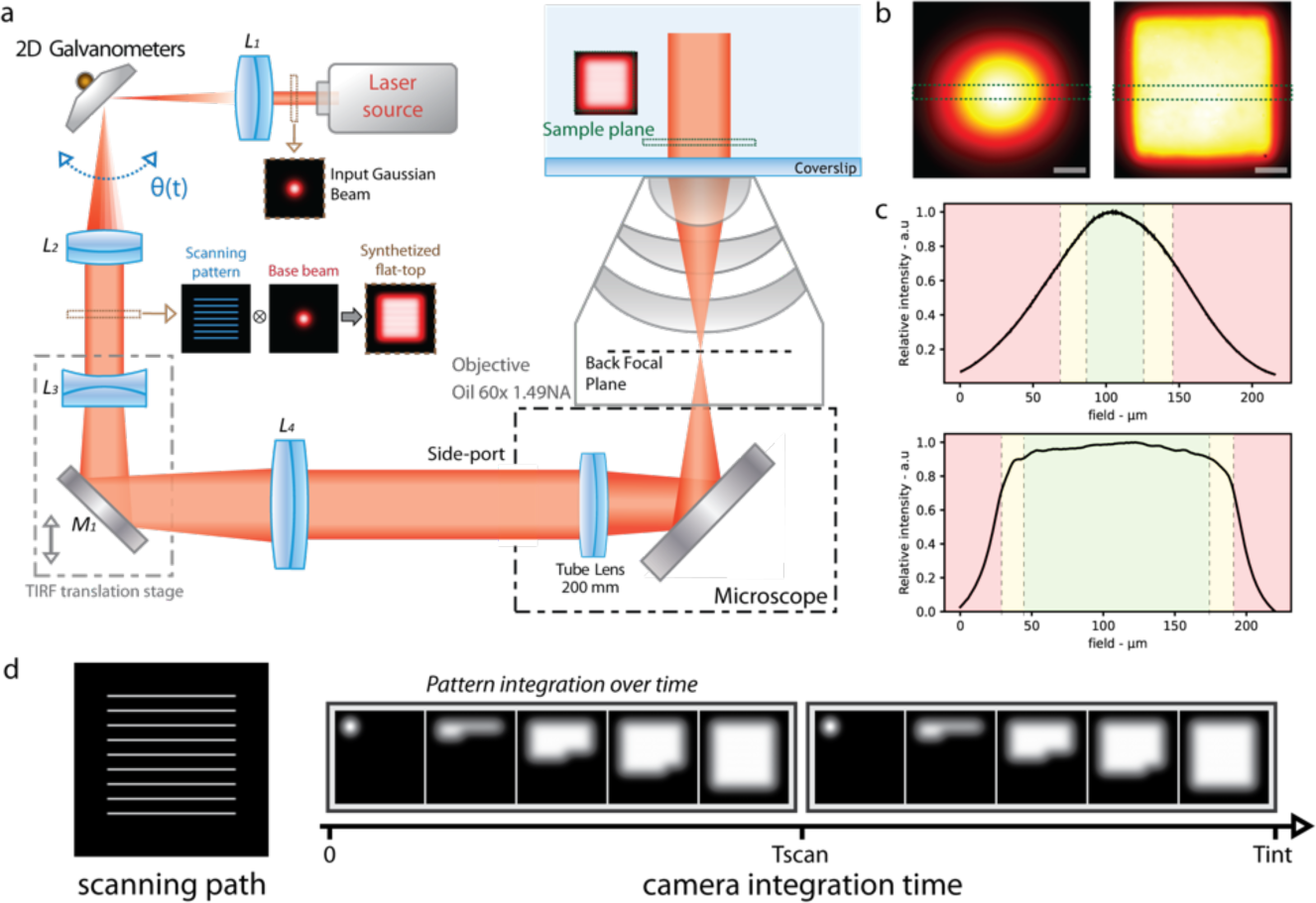
Schematic of ASTER and resulting illumination patterns. (a) Simplified schematic of ASTER setup generating a homogeneous field using a raster scanning pattern. Li are lenses with focal length fi: f1=100, f2=100, f3=-35, f4=250. M1 is a dielectric mirror. A small input Gaussian base beam is scanned in-between the L1 and L2 lenses, resulting in a collimated flat-top profile, which then goes through a TIRF translation stage and is magnified between L3 and L4. After focalization at the BFP of an objective lens, it results in a temporally averaged flat-top excitation profile at the sample. (b) Thin layer of fluorescent Nile-Blue imaged at low laser power with a fixed Gaussian excitation beam (left) and with ASTER (right) raster scanning excitation. Scalebars 40 µm. (c) intensity profiles from (b) of gaussian (up) and ASTER (bottom) illuminations taken along the green dashed area. Dashed lines and colors indicate the field ranges in which intensity is above 90% (green), between 70% and 90% (yellow) or below 70%(red) of its maximum value.(d) Scanning path (left) and generation of the uniform profile over temporal acquisition of the camera. (right). With Tint the camera integration time, the example of a scanning period *Tscan* = *Tint*/2 is shown.

To generate a uniform flat-top excitation over the whole FOV, the scanning needs to meet two criteria. First, the maximum distance between adjacent lines on the beam path has to be lower than 1.7 σ (Supplementary Fig. 1), σ being the standard deviation of the input gaussian excitation beam. Interestingly, decreasing that gap will not affect the flat top so that smaller gaps may be used. For a given field size, this spatial rule defines the minimum number of lines needed to achieve homogeneity (Supplementary Note 2). Second, to avoid stroboscopic effects the flat top must be synthetized under a scanning period *Tscan* that divides the camera integration time *Tint* (Fi. 1d). A typical galvanometer mirror has a repositioning delay of 300 µs, so the number of scanned lines will set the minimal required time to synthesize the flat top profile. Our implementation uses an input excitation beam of σ = 17 µm and gaps between 1.2 - 1.4 σ: ten lines are sufficient to generate a flat-top profile on a 200 µm x 200 µm FOV under 5 ms, which is two times the maximum frame rate of classical sCMOS (100 fps) cameras. In practice, we used camera integration times between 50 to 100 ms and a scanning period of half the integration time so that the flat-top was averaged twice over a single frame, though this number may be modified by adapting the scanning period so that the flat-top is averaged either once, or multiple times (Supplementary Fig. 2). Notably, compared to confocal laser scanning ASTER does not perform point scanning but a continuous scan with a wide input collimated beam and thus can cover large areas much faster.

To characterize the illumination homogeneity and validate our simulations, we imaged a thin layer of fluorescent Nile Blue (Fig. 1b-c). with a classical wide Gaussian beam excitation (σ = 45 µm) and with our ASTER illumination scanning a raster pattern of 150 µm long lines (σ = 17 µm). (Fig. 1c) shows that the ASTER illumination triggers homogeneous Nile Blue fluorescence over a single camera frame, with a square shape matched to typical camera detectors. The resulting flat-top illumination profile is consistent with our simulations and exhibits significant flatness over approximately 130 µm, which could be diminished or increased by adapting the galvanometer input amplitude. In this configuration, if we consider that intensity should remain over 90% of its maximal value for confident quantification over the FOV, the Gaussian illumination would be limited to a 32 µm x 32 µm usable FOV, while ASTER can provide at least a ∼16X larger, 130 µm x 130 µm FOV. On (Fig. 1c) ASTER exhibits gaussian shaped borders that reflects the use of a base gaussian beam of σ=17 µm. A smaller base gaussian beam may be used to sharpen the flat-top borders, but at the cost of slower imaging speed as more lines will have to be scanned. The decrease in brightness at the periphery of the image also stems from vignetting, an effect occurring on all microscope objectives^31^ as light beams emanating from the periphery of the field are partially blocked by optical or mechanical components. We confirm this phenomenon by scanning a large flat-top illuminating the full field of the camera (Supplementary Fig.3). With our 60X magnification and square fields, vignetting is negligible for fields smaller than 160 µm x 160 µm, at 200 µm x 200 µm up to 21% intensity is lost at the corners (affecting 9% of the field), this increases up to 35% loss of intensity on the full 220 µm x 220 µm field of our camera (affecting 18% of the field). In conclusion, even though a wide uniform excitation can be provided, homogeneity is ultimately limited by detection to uniform fields of 160 µm x 160 µm, and relatively uniform fields of 200 µm x 200 µm. By working on a circular field however, a vignetting-free area of 200 x 200 µm^²^ (radius of 113 µm) can be specifically illuminated by scanning an Archimedes spiral (Supplementary Fig. 1).

We then assessed the compatibility of ASTER with inclined, optically-sectioning illumination schemes, where a precise alignment and focusing of the excitation beam in the BFP of the objective is crucial (Supplementary Fig. 4). We focused on TIRF, as it is one of the most common schemes used in SMLM. First, we compared TIRF to epifluorescence illumination (EPI) obtained through ASTER by imaging 3-µm diameter beads, coated with biotin and labelled with AF647-streptavidin (Supplementary Fig. 5) on a 160 µm x 160 µm FOV. As can be assessed on (Fig. 2a-b), due to the spherical shape of the beads the penetration depth of the illumination will be reflected through the beads apparent radii^32,33^. As expected, the measured penetration depth for TIRF excitation is 117 ± 35nm while epifluorescence yields a depth of depth of 865 ± 149 nm, likely defined by the objective’s depth of field (Fig. 2c). The penetration depths for both schemes are uniform over the FOV, and their variations shows no local or global spatial correlation (Supplementary Note 3), demonstrating the absence of a spatial excitation anisotropy. Variation between beads most likely stem from both measurement precision and physical discrepancy of the bead population.

**Fig. 2:**
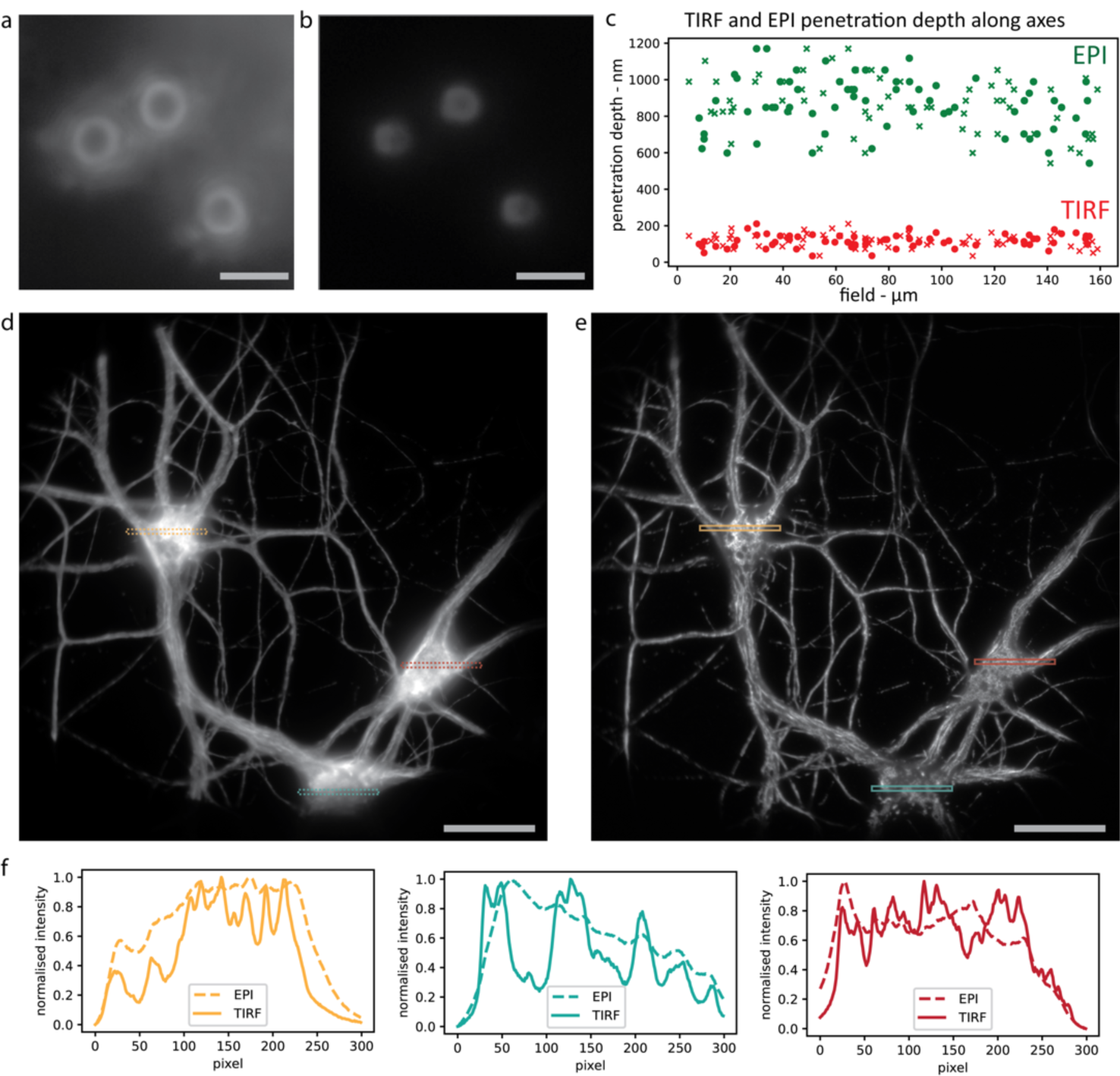
ASTER TIRF Illumination. (a-c) Illumination of 3 µm beads with focus at the coverslip, in ASTER epifluorescence (a) and ASTER TIRF (b) illumination (whole images are shown in Supplementary Fig. 5). Scalebars 4 µm. (c) Measured penetration depth for individual beads in EPI (green) and TIRF (red) illuminations on a large 160µm x 160µm FOV. Cross and circle markers respectively denote measurement along the x and y axis of the sample plane. (d-e) 200 µm x 200 µm imaging FOV of neurons labeled with an anti-β2-spectrin primary and an AF647-coupled secondary antibody, illuminated with raster-scanning ASTER, with a scanning period of 50 ms and an exposure time of 100 ms, in either epifluorescence (d) or TIRF (e) illumination schemes. Scalebars 40 µm. (f) Normalized EPI and TIRF profiles of highlighted areas in (d-e).

To assess the optical sectioning efficiency in experimental conditions, we imaged rat hippocampal neurons labeled for ß2-spectrin, a submembrane scaffold protein lining the neuronal plasma membrane, and revealed with AF647. We compared epi and TIRF configurations on a 200 µm x 200 µm FOV (Fig. 2d-e). The images show that ASTER with TIRF maintains the quality of optical sectioning along the whole FOV: Fluorescence over the cell bodies of neurons (parts that are thicker than the illumination depth) exhibit less blurred fluorescence, and a better signal can be observed compared to the epi-illuminated image, revealing the delicate structure of the neuronal network (Fig. 2f). While a 200ms integration time was used to improve signal to noise ratio, ASTER can provide uniform TIRF excitation under 5ms integration times (Supplementary Fig. 6), which makes it adapted to imaging fast live dynamical processes. A disadvantage of TIRF with classic Gaussian-shaped laser beams are interference speckles: TIRF with ASTER, by contrast, exhibits no such inhomogeneous patterns (Supplementary Fig. 7), as they are likely to be averaged out by beam scanning and camera integration. Even though ASTER is still subject to shadowing effects, it solves both the issues of TIRF interference fringes and non-uniform gaussian illumination, whereas spinning TIRF would prevent further quantification. In conclusion, ASTER leads to TIRF images with a similar quality as spinning azimuthal TIRF, with the added benefit of field uniformity and FOV size versatility. ASTER illumination efficiently provides both uniform spatial illumination and uniform axial optical sectioning for fluorescence microscopy in both epi and TIRF illumination.

### Large Field Uniform SMLM Imaging

Next, we applied ASTER to SMLM experiments, namely DNA-Point Accumulation in Nanoscale Topography (PAINT) and STochastic Optical Reconstruction Microscopy (STORM). To assess the effect of ASTER illumination FOV size and homogeneity in SMLM experiments, and compare it to a classical Gaussian illumination, we first imaged three-spot, 40 nm spaced nanorulers using DNA-PAINT (Fig. 3). Three different TIRF excitation schemes (Gaussian, σ =45 µm), ASTER on a 70 µm x 70 µm FOV, and ASTER on 120 µm x 120 µm FOV) were used, with the other parameters remaining identical. In the single molecule regime, each of the three nanoruler spot acts as a source of blinking fluorescence, resulting in a set of localizations spread by the pointing accuracy of each blinking event. For analysis we applied the following algorithm: first, individual nanorulers were isolated by DBscan clustering^34^, then for each individual nanoruler the point cloud corresponding to the three spots was fitted by a Gaussian Mixture Model (GMM) assuming three normal distributions. The GMM assessed the most probable mean position and standard deviation of each spot (see Methods, Supplementary Fig. 8). The mean standard deviation of all spots was then considered as the experimental localization precision for each individual nanoruler.

**Fig. 3:**
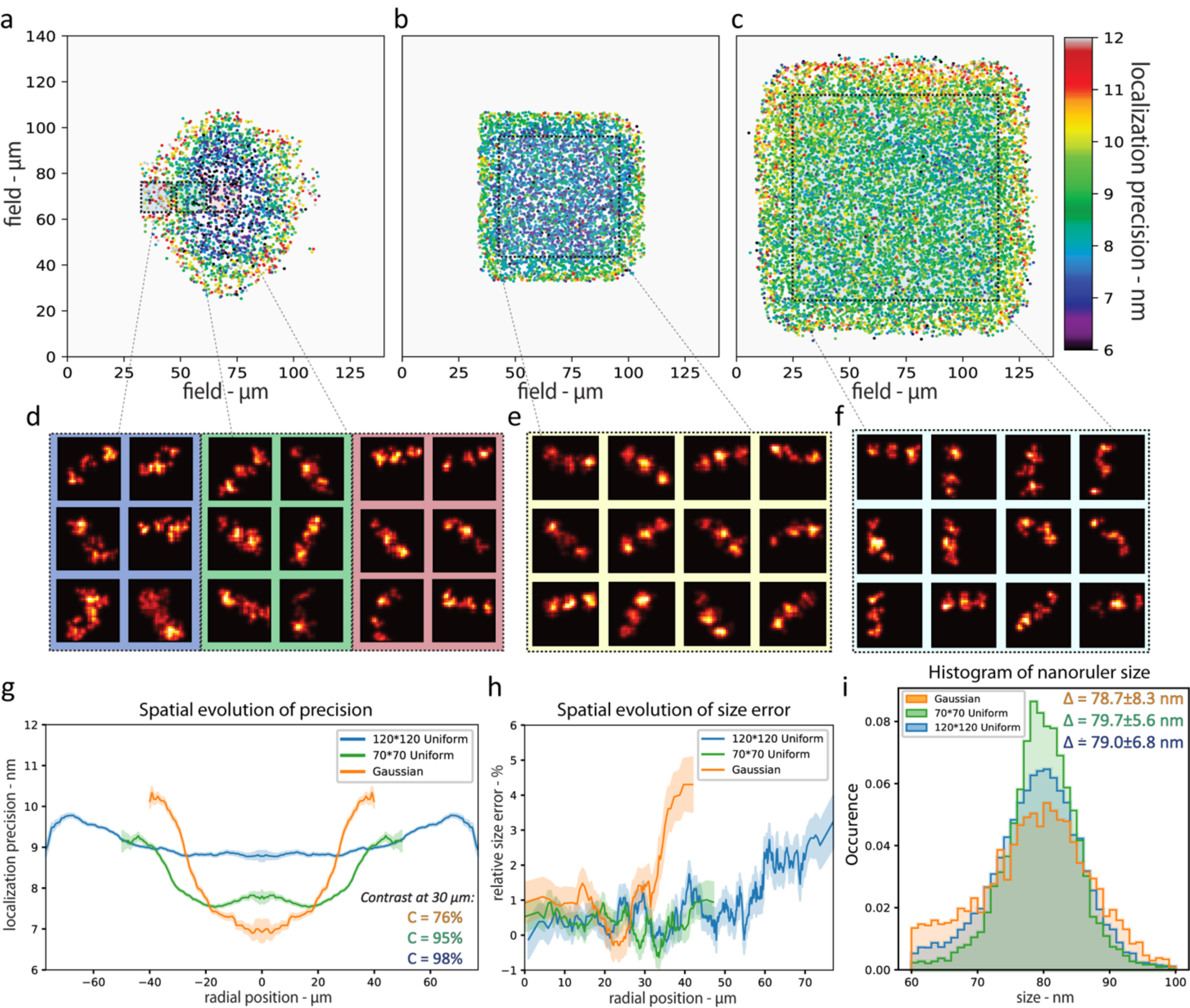
Nanorulers imaging for localization precision estimation. (a-f) DNA-PAINT imaging of 40 nm spaced 3-spots nanorulers, obtained with Gaussian (left), ASTER small field of view (70 µm x 70 µm, middle), and ASTER large field of view (120 µm x 120 µm, right) illuminations. (a-c) are resulting localization precision maps where each point represents the average precision for one individual nanoruler (3 spots). (d-f) are nanoruler superresolution images (5 nm pixel size), taken randomly from highlighted areas in (a-c). (g) Measured localization precision along FOV radius for each excitation scheme (symmetrized). For each colored curve, the surrounding transparent curve indicates the standard deviation around the mean precision at a given radius. (h) Resulting size estimation error along FOV radius for each excitation scheme. A size error above 0 indicates that the nanoruler spots were measured less than 80 nm apart. Each colored curve is surrounded by a transparent curve that indicates the standard deviation around the mean size. (i) Resulting size measurement histogram for each excitation scheme. The mean and the standard deviation are indicated in the upper right corner.

The Gaussian excitation resulted in a bell-shaped localization precision map (Fig. 3a,g).: at the center of the FOV, the localization precision is only 7 nm, but it quickly increases with the distance to FOV center. At 20 µm it is 8 nm, and up to 11 nm at the edges of the FOV, 1.6 times worse than at center (Fig. 3d,g). Meanwhile, ASTER excitation on a similar FOV provided a localization precision of 7.9 nm ± 0.9 nm - ranging from 7.5 to 8 nm at 30 µm from the center of the field (Fig. 3b, g). On a large 120 µm x 120 µm FOV with similar parameters, ASTER provided a 9.2 nm ± 1.1 nm localization precision (Fig. 3c, g), from 8.8 at the center of the field up to 9.5 nm at a 60µm radial distance. This means that a 20X increase in the FOV size came at the cost of a 1.2 worse localization precision. It is conceivable that a localization precision below 9 nm could be reached by carefully optimizing imaging parameters such as laser power, optical sectioning and camera integration time. Moreover, the inhomogeneity of the Gaussian illumination impacted the size estimation of the nanorulers along the FOV (Fig. 3h). We measured the end-to-end size of the identified nanorulers (ground truth value 80 nm). Gaussian beam illumination images yield a fairly constant relative size error of 1% at the center of FOV, rising to 3% at 35 µm from the center. ASTER provided homogeneous measurements on a wider FOV: the relative size error remained at 0.5% up to 45 µm from the center of the FOV. All the illumination conditions resulted in similar mean values for the nanoruler size (Fig. 3i), but we observed an increased number of cases where the size was underestimated to 60-70 nm with the Gaussian beam illumination, indicating a poor single molecule regime.

We then turned to STORM experiments on biological samples. Traditionally, STORM demands strong laser power (>2kW/cm^²^) to drive organic fluorophores into a blinking regime^35,36^. To induce a satisfactory blinking regime on a 200 µm x 200 µm FOV thus requires the use of 1-5 W power lasers^27^. However, as ASTER provides locally high excitation irradiance on a short time scale, and a lower global average excitation on longer scales, it may partly overcome this irradiance threshold rule. We applied ASTER in HiLo to a direct STORM experiment and found that even with reasonable laser power (<0.3 W at BFP), ASTER was able to induce and maintain a densely-labeled sample in the sparse single molecule regime (<1 molecule per µm^3^) on large FOVs (Fig. 4) where conventional illumination would fail. It appears that the high but intermittent local excitation intensity (∼12kW/cm^²^) nonetheless sends most of the molecules in a long-lived dark state efficiently, as is expected for high irradiances^37^.

**Fig. 4:**
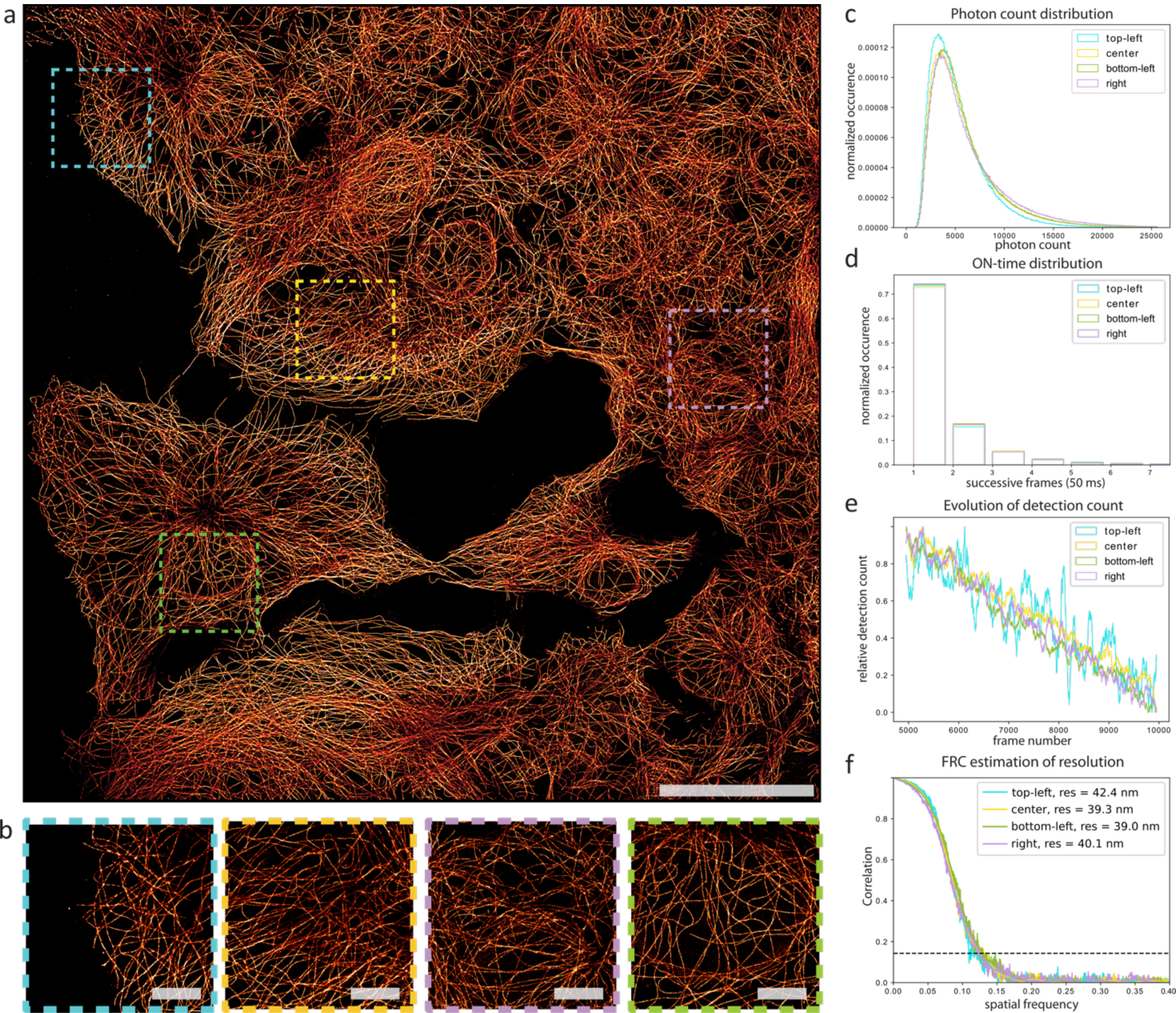
STORM imaging using ASTER. (a) ASTER STORM imaging of COS-7 cells labeled for microtubules and an AF647-coupled secondary antibody, FOV size 200 µm x 200 µm, 20,000 frames at 20 fps. Excitation consisted in a ten-line scan with a laser power of 250 mW at the BFP, a gap of 1.4σ and a 25 ms scanning period. Scalebar 50 µm. (b) Zoomed views of highlighted areas in (a). Scalebar 10 µm. (c) Photon count distribution histogram for highlighted areas in (a). (d) Blinking ON-time distribution for highlighted areas in (a), expressed in number of successive frames (50 ms camera integration time). (e) Temporal evolution of detection count for highlighted areas in (a). (f) FRC estimation of resolution for highlighted areas in (a).

(Fig. 4a) shows a SMLM image of a COS cell labeled for microtubules and an AF647-coupled secondary Fab_2_ antibody. It was obtained using 20,000 frames and a 50 ms camera integration time, using ASTER with 0.3 W laser power, raster scanning a 200 µm x 200 µm FOV (ten lines). The microtubules are well resolved throughout the whole FOV; the zoomed images in (Fig. 4b) show the image quality in several different parts of the image. Analysis of these regions revealed comparable photon distributions, blinking ON-time and localization density during acquisition (Fig. 4c-e). Even though a slight decrease in photon count can be noticed at the edge, other blinking characteristics remain unchanged and suggest once again inhomogeneous detection due to vignetting at the periphery of the objective. Analysis from regions of an image acquired with a classical Gaussian illumination showed significant differences between the regions, underlining the detrimental effect of in-homogeneous Gaussian illumination (Supplementary Fig. 9a-f). We assessed the experimental image resolution with Fourier Ring Correlation (FRC) analysis^38^ (Fig. 4f), and found that the region subject to vignetting had a close resolution (42 nm) to the other areas (39 – 40 nm) indicating that vignetting does not significantly impact the uniformity on our FOV. This confirms ASTER’s ability to obtain uniform blinking and resolution on large 200 µm x 200 µm FOVs in STORM. Noteworthy, we do not notice artefact emerging from the temporal scanning of the beam, indicating that the position of a fluorophore relative to the scanning part does not matter. Further experiments (Supplementary Fig. 10) suggest that all STORM blinking properties remain similar as long as the same mean irradiance is provided.

ASTER thus is compatible with both DNA-PAINT and STORM experiments even with typical lasers currently used on SMLM microscopes with output power below 1W. In SMLM, because of a required pixel imaging size around 100nm, camera chip finite size will ultimately limit the FOV, the largest uniform FOV reported so far being 221 µm x 221 µm by Zhao et al^24^. However, their implementation did not perform TIRF and required multiple lasers with >1W output power plus a vibration motor to reduce speckles. In all cases, imperfections from the detection path will limit the maximum achievable FOV. To overcome this limit, we stitched four 150 µm x 150 µm uniform STORM images, resulting in a 300 µm x 300 µm image (Supplementary Fig. 11a-b) with minimal overlap and high uniformity. Stitching results in minimal artefacts in the overlapping edge areas, but slightly suffers from temporal effects on photon count and molecule density, mostly due to buffer consumption between acquisitions (Supplementary Fig. 11c). To limit temporal effects, one may choose to speed up STORM experiment by increasing the global irradiance^39,37^. With ASTER, this can be done by reducing the amplitude of scanning. We scanned five 25 µm-long lines in 5ms to reach an effective irradiance of 27 kW/cm^²^, this allowed to perform fast STORM imaging of microtubule in under 100 seconds on a classical FOV (Supplementary Fig. 12). Such experiment is less prone to drift and highlights the practical versatility of ASTER for optimizing STORM experimental needs^40^.

As ASTER homogenously illuminates large FOVs, it extends the possibility of quantitative analysis of nanoscopic structures to whole cells or group of cells. To obtain a precise view of a biological structure at the nanoscale, it is crucial to leverage the imaging of a large number of similar structures. This allows to not only obtain their average characteristics, but also the individual variation of these characteristics caused by biological variability. We imaged clathrin clusters and clathrin-coated pits by STORM in COS-7 cells (Fig. 5a-c) and applied a cluster analysis. Three COS-7 cells were imaged at once over a large 140 µm x 140 µm FOV, containing approximately 20,000 individual clathrin clusters. In comparison, a classical 30 µm x 30 µm FOV would have yielded ∼1,500 pits. The high number of clathrin clusters identified on resolution-uniform images allowed for population estimation from the characteristics of clusters. We picked specific parameters such as diameter and hollowness (see Methods). We were able to distinguish four populations from the cluster diameter distribution, as fitted with normal distributions (Fig. 5b). Small-diameter clusters (below 80 nm, blue and green population of Fig. 5b) likely correspond to pits in formation, while large ones (orange and red populations in Fig. 5b) are likely to be fully assembled pits. We specifically extracted large, hollow clathrin assemblies based on the ratio between the diameter and the spatial dispersion of fluorophores. Hollow clathrin assemblies with diameters of 80-200 nm would be of typical size for the large clathrin-coated pits found in fibroblasts^41^. Interestingly, some large, hollow pits showed more than one fluorescence “holes” within them, suggesting that they are either assemblies of smaller pits or that the fenestration of clathrin cages^42^ (pentagon or hexagons of 18 nm side length) can sometimes be resolved.

**Fig. 5:**
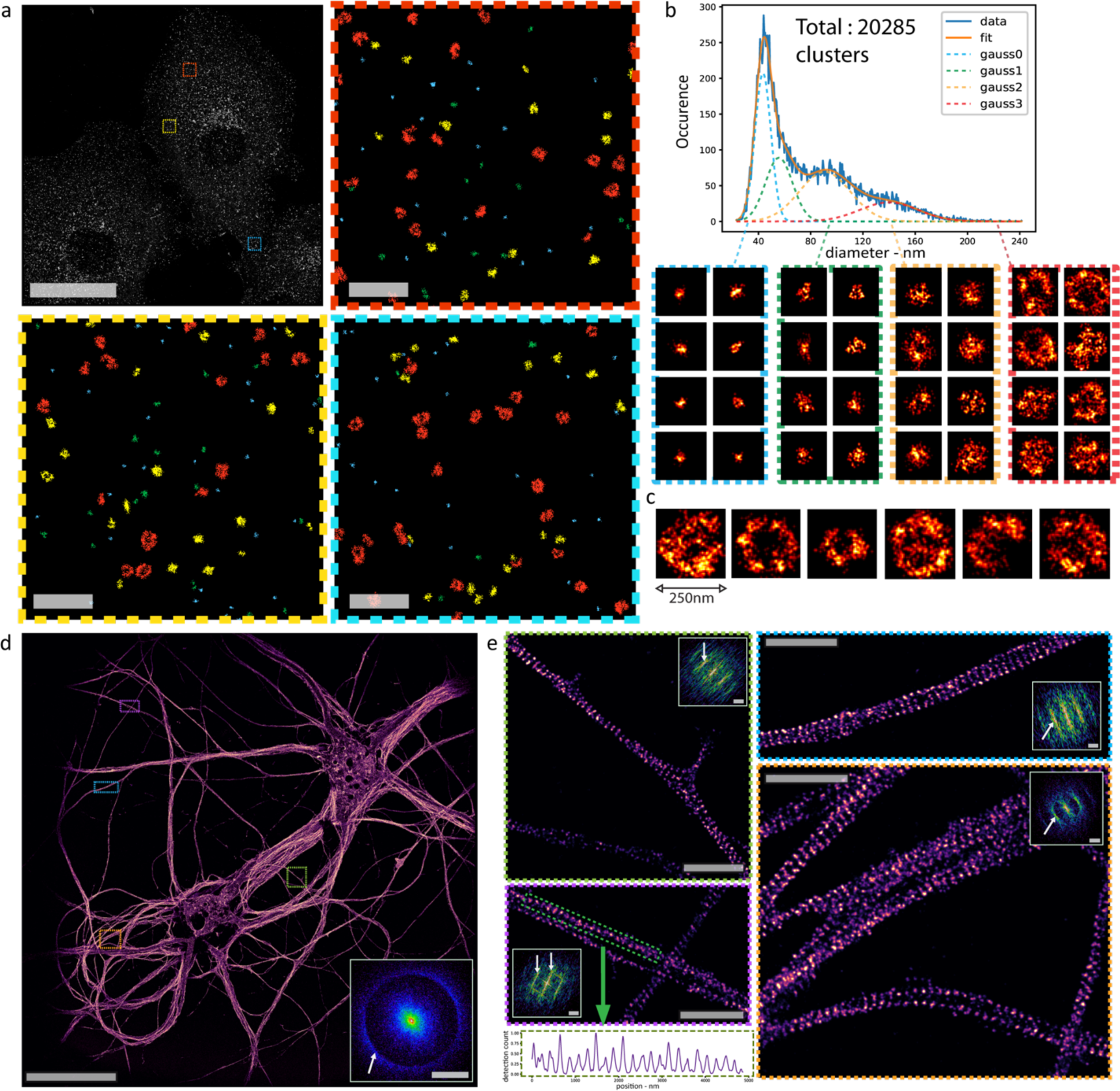
ASTER applications for single molecule localization microscopy. **(a-c)** ASTER STORM imaging and cluster analysis of COS-7 cells labeled for clathrin heavy-chain and an AF647-coupled secondary antibody. (a) Final 140 µm x 140 µm image (top left) and close-up views of the highlighted regions (colors encode cluster affiliation). Pixel size is 10 nm. (b) shows the distribution of the diameter of clathrin clusters and highlights four potential populations, that can be fitted with gaussian functions. Below are 5 nm-pixel images of individual clathrin related clusters, each area corresponding to a specific population. (c) shows images (5 nm pixel size) of large, hollow clathrin clusters likely corresponding to large clathrin-coated pits. **(d**,**e)** ASTER STORM imaging and structural analysis of neurons labeled for β2-spectrin and AF647-coupled secondary antibody. (d) 200 µm x 200 µm STORM image obtained with ASTER (30 nm pixel size). The two-dimensional Fourier transformation (inlet) exhibits a circular frequency pattern corresponding to a ∼190 nm periodicity of the staining that is present along all axons. (e) Zoomed views of regions in (d) revealing the periodic cytoskeleton along single axons (pixel size is 10 nm). Insets show their respective two-dimensional Fourier transformation, showing line patterns at corresponding to the known 190 nm periodicity of the axonal spectrin scaffold. bottom left image shows the intensity profile along the highlighted green line, revealing the same the 190 nm periodicity.

The large FOV provided by ASTER illumination coupled with large-chip sCMOS cameras also have interesting application for imaging neuronal cells, which grow axons over hundreds of microns in culture. Traditionally, SMLM imaging of axons has been limited to <50 µm segments of axons, impeding the visualization of rare structures and the definition of their large-scale organization^43,44^. We labeled rat hippocampal neurons for β2-spectrin, a protein that forms a periodic sub membrane scaffold along axons by linking actin rings^45–47^. A 200 µm x 200 µm FOV allowed visualizing the dendrites and cell body of two neurons, and a large number of long axonal segments (Fig. 5d, Supplementary Fig. 13). The zoomed views confirm the quality and resolution of the resulting image: the periodic 190 nm organization of axonal spectrin is clearly visible, as confirmed by the corresponding Fourier transform of the images. The Fourier transform of the whole image exhibits a sharp ring at the corresponding frequency, because the banded pattern of β2-spectrins appears in axons running in all directions. On the zoomed images, the β2-spectrin along axons in one direction results in a direction-dependent frequency band on the Fourier transform, corresponding to the 190 nm spacing.

## Discussion

We implemented and characterized ASTER, a hybrid scanning and wide-field illumination technique for optimized wide-field fluorescence microscopy and Single Molecule Localization Microscopy (SMLM) over large fields of view (FOV). ASTER generates uniform excitation over a tunable FOV without limiting acquisition speed. It has advantages over state-of-the-art uniform illumination schemes by its efficiency, flexibility and ability to perform uniform optical sectioning schemes, such as HiLo and TIRF illuminations. We demonstrate TIRF imaging on rat hippocampal neurons on 200 µm x 200 µm, the maximal uniform FOV achievable with our x60 magnification objective. In PAINT, we demonstrate a uniform localization precision over large FOVs (9.2 ± 1.1 nm over 120 µm x 120 µm), and even better resolution on small FOVs (7.9 ± 0.9 nm over 70 µm x 70 µm).

ASTER also proved to be an efficient excitation method for STORM imaging experiments. Against common belief that STORM requires a strong continuous irradiance (∼2 kW/cm^2^), ASTER induced uniform blinking dynamics at lower mean irradiance (< 0.5 kW/cm^2^), but with a high instantaneous irradiance (∼12 kW/cm^²^) over a large 200 µm x 200 µm FOV, alleviating the need for expensive and dangerous high-power lasers. We present biological applications by directly imaging the periodic 190 nm organization of axonal spectrin on several long axon segments from neurons. By imaging clathrin-coated pits in multiple COS-7 cells in one acquisition, we increase the number of identified clusters by a factor of 20 compared to the typical FOV of a STORM acquisition and enhance statistical analysis.

ASTER can be combined with stitching schemes, alternative objectives, adaptive detection setups and camera chips to cover even wider FOV. Combination of ASTER with the ASOM^48,49^ could be particularly interesting as hyper large fields may be imaged without moving the sample. In SMLM, the field is regularly limited to a maximum of 200 µm x 200 µm, however in classical widefield microscopy ASTER can be used with smaller magnification objectives to image larger FOVs and would be a great choice for imaging structures on larger scales. We conclude that ASTER represents a versatile and innovative tool, especially suited for SMLM. It exhibits robust uniformity and reliability, as well as adaptability to variable FOV sizes. ASTER may be used in combination with improved detection schemes, such as multicolor imaging or strategies to encode axial information^50,51^ (see Supplementary Fig. 14). It could be beneficial to setups that modify the excitation to enhance resolution by counting photons ^52–55^ but ASTER implementation in such case would be challenging. The resulting uniformity may be used for demultiplexing or stoichiometry experiments^56,57^, as well as in buffer characterization and other fields such as photolitography^58,59^. Finally, ASTER has potential applications in non-uniform excitation schemes, such as using smaller beams to concentrate power^60^, exciting specific areas of a sample^61^ or creating a patterned irradiance on complex samples by using adaptive scanning strategies.

## Acknowledgements

We acknowledge people who provided valuable advice and support in the course of this study; particularly P. Jouchet and C. Cabriel. We thank S. Vassilopoulos for discussions about clathrin organization in cells. M. Bardou and F. Boroni-Rueda helped with cell culture and labelling. C. Hubert and E. Fort provided the lasers and galvanometer scanners.

## Author contributions

A.M., N.B. and S.L.-F. conceived the project. All authors contributed to the writing and editing of the manuscript. K.F and C.L provided neuronal samples. Experimental acquisitions and analyses were performed by A.M.

## Methods

### Optical setup

We used a Nikon Eclipse Ti inverted microscope with a Nikon Perfect Focus System. The excitation was performed with a ELERA laser (638 nm) from ERROL. 6215H galvanometers from Cambridge Technology were controlled with a RIGOL DG5252 waveform generator. To maintain telecentricity, all distances between lenses are equal to the sum of their respective focal lengths. Both excitation and detection went through the left camera port of the microscope to prevent undesired cropping. To this end, the dichroic is put in front of the side port and reflects the excitation beam (not shown in Fig. 1).Fluorescence was collected through an Olympus x60 1.49NA oil immersion objective, a relay-system, and recorded on a 2048*2048 pixel sCMOS camera (Orca-Flash 4 v3, Hamamatsu). The optical pixel size was approximately 108 nm.

### Calibration sample preparation and imaging

#### Beads

Beads are 3 µm radius biotin-polystyrene microspheres (Kisker Biotech,PC-B-3.0) on which we attached Alexa Fluor (AF) 647 functionalized with streptavidin (Life Technologies, S21374). We prepared a solution containing 500 µL of water, 500 µL of PBS, 35 µL of microsphere solution, and 0.34 µL of AF647. This solution was centrifuged 20 min at 13.4 krpm. The liquid was then removed and replaced with 100 µL of PBS, followed by 5 minutes vortexing to dissolve the deposit. 50 µL of the final solution was then pipetted on to a glass coverslip and left for 20 min so that beads would have time to deposit. Finally, we added 500 µL of imaging dSTORM buffer (dSTORM smart kit, Abbelight). Images were taken at low laser power and integrated over 100 ms, for an ASTER scan period of 50 ms.

#### Nanorulers

Nanorulers (Gattaquant, PAINT-40R) consist in three aligned spots, separated by 40 nm and are labelled with ATTO655 fluorophores. To switch from ASTER to a gaussian illumination, a constant offset was applied to galvanometers and a beam magnifier was placed between the laser and galvanometers. Parameters for imaging were chosen while optimising the blinking with the wide-gaussian excitation (σ = 45 µm): laser power of 200 mW, fixed TIRF configuration and an integration time of 100 ms; parameters were maintained for each acquisition. During the acquisition on the 120 µm x 120 µm FOV, the blinking of fluorophores was slow, so fluorophores generally appeared on subsequent frames. and required larger integration times or post-processing to merge them. This became apparent in the loss in resolution.

### Fluorescence immunolabeling

#### Neuronal culture

Rat hippocampal neurons in culture were prepared according to the Banker protocol^62^. Briefly, E18 Wistar rat embryo hippocampi (Janvier labs) were dissected, then cells were homogenized and plated in B27-containing Neurobasal medium on Poly-L-Lysine treated #1.5H glass coverslips (Marienfeld, VWR) to a density of 4000 cells per cm^2^. The neurons were then co-cultured with glia cells - neuron coverslip upside down, separated from the glia on the bottom of the petri dish by wax beads. Mature neurons were fixed after 14 days in culture. All procedures followed the guidelines from European Animal Care and Use Committee (86/609/CEE) and were approved by local ethics committee (agreement D13-055-8).

Immunolabeling of neurons was performed as described recently for optimized SMLM sample preparation^47,63^. Neurons were fixed using 4% paraformaldehyde (Delta Microscopie, #15714) and 4% (w/v) sucrose in PEM buffer (80 mM PIPES, 2 mM MgCl_2_, 5 mM EGTA, pH 6.8) for 20 minutes at RT. Cells were then rinsed with 0.1 M phosphate buffer. Blocking and permeabilization were performed in ICC buffer (0.2% (v/v) gelatin, 0.1% Triton X-100 in phosphate buffer) for 2 hours on a rocking table. Primary antibodies diluted in ICC were incubated overnight at 4°C, rinsed and incubated with the secondary antibodies diluted in ICC for one hour at room temperature. After a final rinse with ICC and phosphate buffer, the samples were stored in phosphate buffer with 0.02 % (w/v) sodium azide before imaging. For immunolabeling, we used mouse anti β2-spectrin (BD Sciences, #612563, 2.5 µg/ml) and donkey anti-mouse AF647 (ThermoFisher, #A31571, 6.67 µg/ml).

#### Cell line culture

COS-7 cells were grown in DMEM with 10% FBS, 1% L-glutamin and 1% penicillin / streptomycin (Life Technologies) at 37°C and 5% CO2 in a cell culture incubator. Two days later, they were plated at medium confluence on cleaned, round 25 mm diameter high resolution 1.5’’ glass coverslips (Marienfield, VWR). After 24 hours, the cells were washed three times with PHEM solution (60 mM PIPES, 25 mM HEPES, 5 mM EGTA and 2 mM Mg acetate adjusted to pH 6.9 with 1 M KOH). For preparation of STORM microtubule imaging, we added an extraction solution (0.25% Triton, 0.025% Glutaraldehyde in PEM) for 30 s then a fixation solution (0.5% glutaraldehyde, 0.5% Triton in PEM) for 12 min followed by a reduction solution (NaBH_4_: 0,1 % in PBS 1X) for 7 minutes. For clathrin we directly fixed with a 4% PFA solution. Extraction and fixation solutions were pre-warmed at 37°C. Cells were then washed 3 times in PBS before being blocked for 15 min in PBS + 1% BSA + 0.1% Triton. Labelling was performed in a similar solution with intermediary washing steps. α-tubulin (Sigma Aldrich, T6199) and clathrin heavy-chain (Abcam, ab2731) primary antibodies were conjugated with Rb-AF647 (Life Technologies, A21237). Cells were finally post-fixed for 16 minutes in 3.7% Formaldehyde and reduced for 10 min with NH_4_Cl (3mg/mL).

### Biological sample imaging

#### Wide-field fluorescence imaging

TIRF imaging on neuronal sample was done at 200 ms integration times and a low 30 mW laser power. Samples consisted in β2-spectrin labelled with AF647. Output angle was adjusted with a translation stage^64^ until penetration depth was roughly around 200nm.

#### STORM imaging

STORM imaging on COS-7 cells (microtubules and clathrin) and neurons (ß2-spectrin) was performed at 50ms exposure time using a HiLo illumination configuration. A STORM buffer (Abbelight Smart kit) was used to induce most of the molecules in a dark state. The sample was lit with laser powers of approximately 250 mW in the objective BFP and a scanned gaussian beam of σ=17 µm. Except for clathrin, all data acquisition was excited with an ASTER excitation scanning ten lines in 25 ms and a gap of 1.4σ. For STORM on clathrin, the labeling was dense and blinking was slightly optimized by reducing the excited FOV to 140 µm x 140 µm, scanning eight lines in 25ms with a gap of 1.2σ. The acquisitions were performed and analyzed using the Nemo software (Abbelight). Localization consisted in a wavelet segmentation after median background removal, followed by gaussian fits of individual point spread functions. Sample was drift-corrected using a classical redundant cross-correlation algorithm.

#### Image acquisition, processing and analysis

Neo-Live (Abbelight) was used for image acquisition. Single molecule analysis was performed either with a home-made Python 3.7 code or with Neo-Analysis (Abbelight).

Data processing such as analysis and measurement of beads radius from (Fig. 2), nanoruler from (Fig. 3) and clathrin from (Fig. 5) was performed in Python; whose code is available online.

#### Beads

Beads (microspheres) were detected on the TIRF image: we first applied a Laplace filter from SciPy library, followed by low-pass filtering in Fourier space to diminish noise. Use of an intensity threshold then proved sufficient to efficiently detect individual beads. Peaks positions were measured via local extrema algorithms.

#### Nanorulers

Nanorulers analysis focused on resulting X,Y coordinates. A preliminary DBscan clustering was used to localize and filter out lonesome localizations. Then a more precise DBscan was used to distinguish individual groups of three-spots and associate a number to each of them. DBscan typically consider core points, which are point with at least *mpts* neighbors in a surrounding *epsilon* radius, then iteratively add adjacent points. Parameters of this secondary scan were: epsilon=50 nm and minimum number of points mpts=10. With these parameters, adjacent spots belonging to a similar nanoruler array were grouped together, while unwanted associations of adjacent nanorulers were minimized. For each group, a Gaussian Mixture Model (GMM) clustering was used to estimate parameters from three gaussian distribution. GMM also estimates the mean and standard deviation of each spot, which allowed for size estimation and localization precision measurements. Nanorulers with too few points or extraordinary distance estimations were thrown away.

#### Clathrin clusters

Clathrin analysis was primarily performed via a DBscan clustering, with an epsilon parameter of 35 nm and a minimum number of points of 25. This clustering method localized each individual cluster of close points. For each of these cluster we calculated several parameters, such as the mean position and the effective diameter, Feret’s diameter^65^, the hollowness, the angle of orientation and eccentricity. Effective diameter and mean position were calculated by minimizing radial dispersion among points. Hollowness consisted in the ratio between the mean radius value, divided by the standard deviation of radius, and was found to be rather independent of the size of the cluster. Among all parameters, the diameter and the hollowness proved to be the most relevant in term of describing cluster distributions.

#### Neurons

Fourier transform was performed on 2D histogram images via the fft2 function from the numpy.fft library of Python.

## Data availability

SMLM large data files (>20Go) are available from the corresponding author on reasonable request. Other data files are available on Zenodo (DOI: 10.5281/zenodo.3814322) as well as related analysis code.

## Code availability

Code is available online on GitHub at the following link: https://github.com/AdrienMau/ASTER_code and Zenodo (DOI: 10.5281/zenodo.3814322).

## Supplementary Figures

**Supplementary Figure 1:**

Field synthesis with different gaps between scanning lines and different scanning patterns

**Supplementary Figure 2:**

Chronogram of ASTER scanning excitation and camera integration.

**Supplementary Figure 3:**

Vignetting limits the maximum homogeneous FOV

**Supplementary Figure 4:**

Implementation of TIRF and oblique illumination in classical and ASTER excitation schemes

**Supplementary Figure 5:**

Beads for calibrations of optical sectioning

**Supplementary Figure 6:**

Full-field TIRF uniform ASTER excitation at 5ms integration time.

**Supplementary Figure 7:**

Homogeneity of ASTER TIRF excitation compared to azimuthal spinning TIRF

**Supplementary Figure 8:**

Workflow analysis of nanoruler images

**Supplementary Figure 9:**

Gaussian illumination effects in single molecule STORM microscopy

**Supplementary Figure 10:**

Impact of scanning on STORM blinking properties

**Supplementary Figure 11:**

Stitching of STORM images resulting in a 300 x 300 μm^²^ field of view

**Supplementary Figure 12:**

Fast STORM experiment acquired at 5ms integration time

**Supplementary Figure 13:**

STORM 200 µm x 200 µm image of neuronal β2-spectrin.

**Supplementary Figure 14:**

STORM 200 µm x 200 µm 3D image of COS-7 cells labeled for microtubules

**Supplementary Notes 1:** Comparison of uniform excitation methods

**Supplementary Notes 2:** Relation between minimum frame rate and field size with ASTER

**Supplementary Notes 3:** Uncertainties in measurement of microbead excitation depth

**Supplementary Figure 1:**
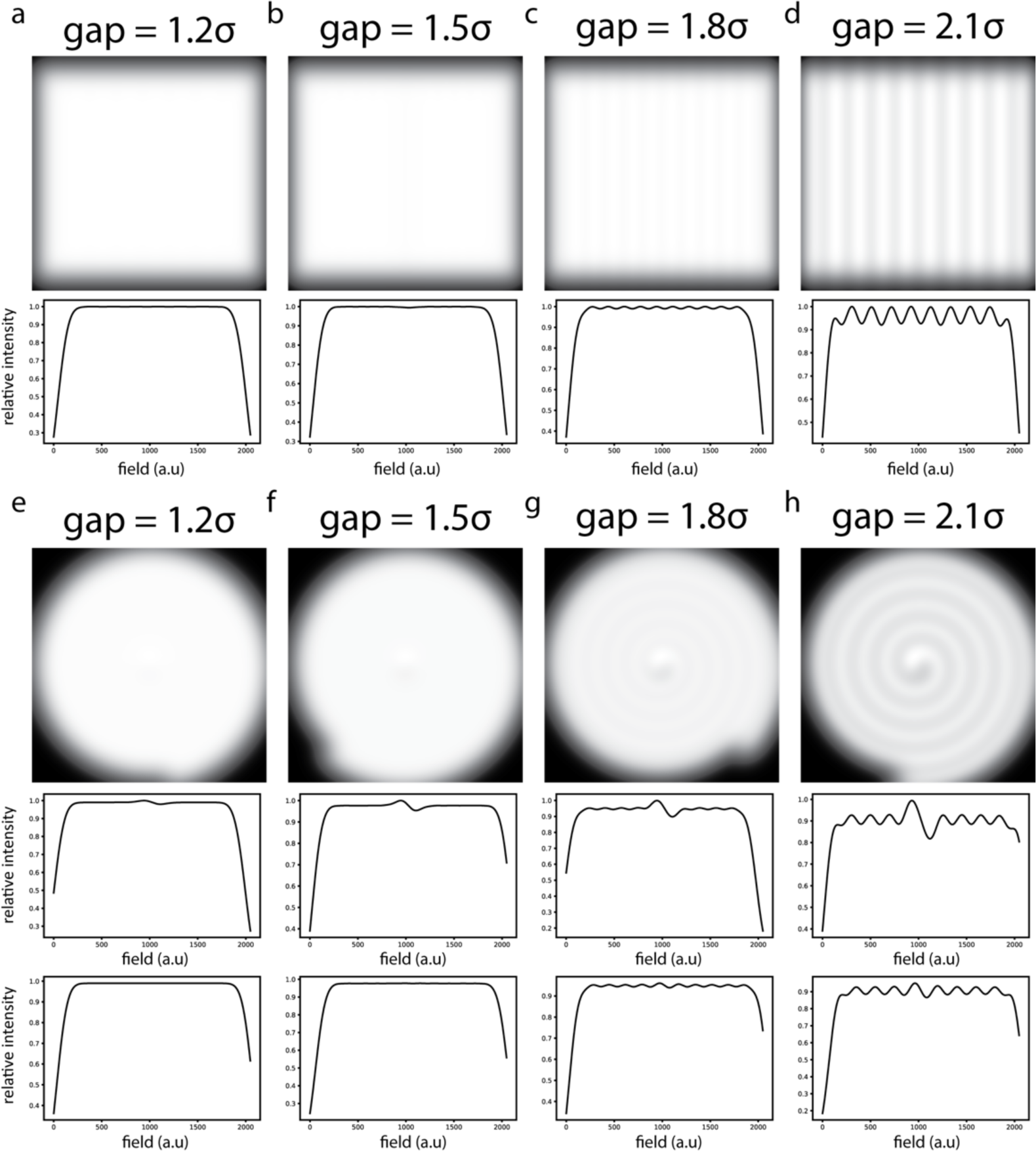
Field synthesis with different gaps between scanning lines and different scanning patterns. (a-d) Resulting illumination for a raster-scanning pattern and different line gaps. Under each image, resulting horizontal profiles taken at center are shown. (e-f) Resulting illumination for scanning an Archimedes spiral at different line gaps. Under each image, resulting illumination profiles along vertical (up) and horizontal (bottom) axes are shown. σ denotes the standard deviation of the scanned gaussian beam.

**Supplementary Figure 2:**
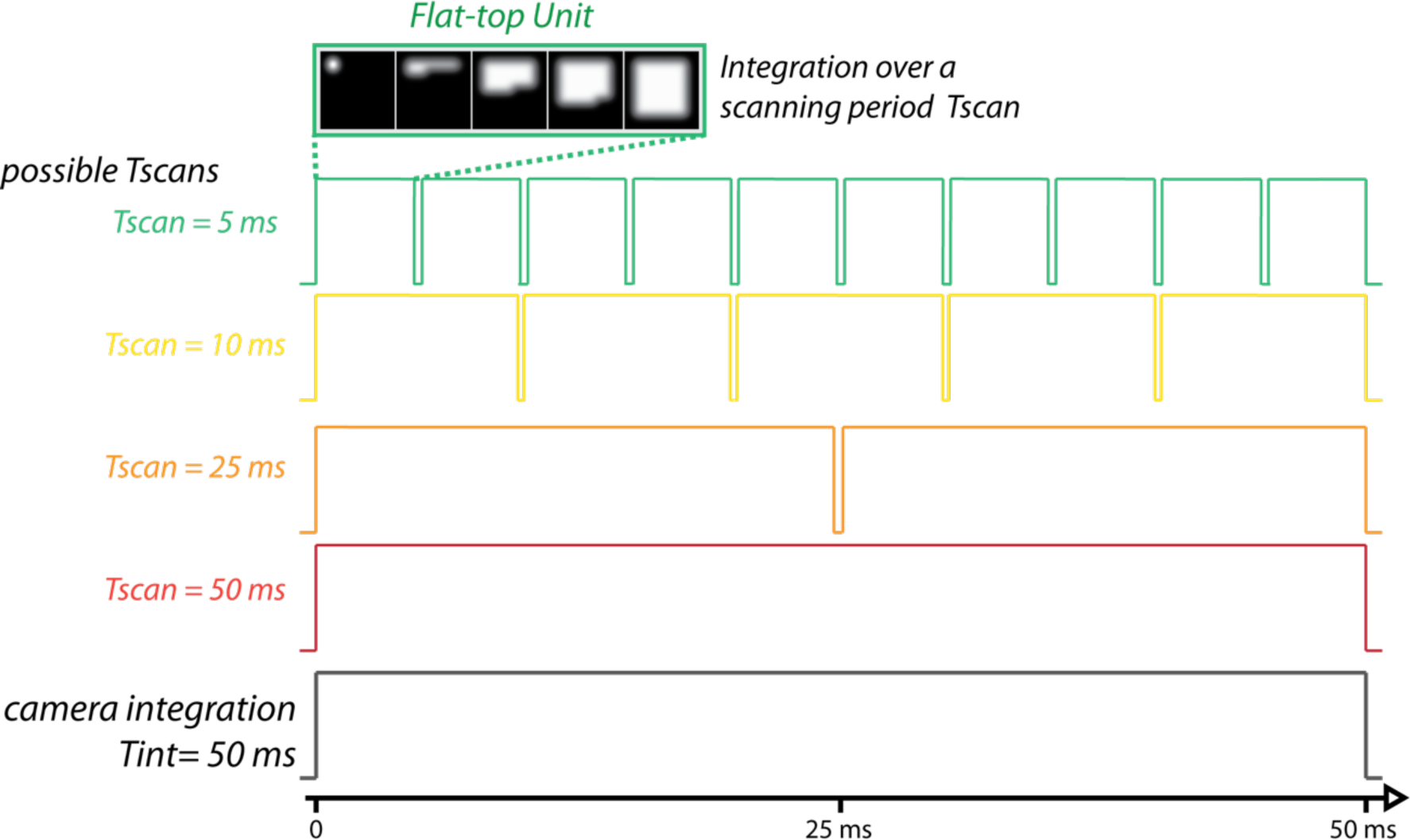
Chronogram of ASTER scanning excitation and camera integration. For a given camera integration time (here Tint=50 ms), the scanning period *Tscan* of ASTER should divide Tint so that a finite number of flat-tops are generated over the integration. Examples for *Tscan* values are shown, namely 5 ms, 10 ms, 25 ms and 50 ms.

**Supplementary Figure 3:**
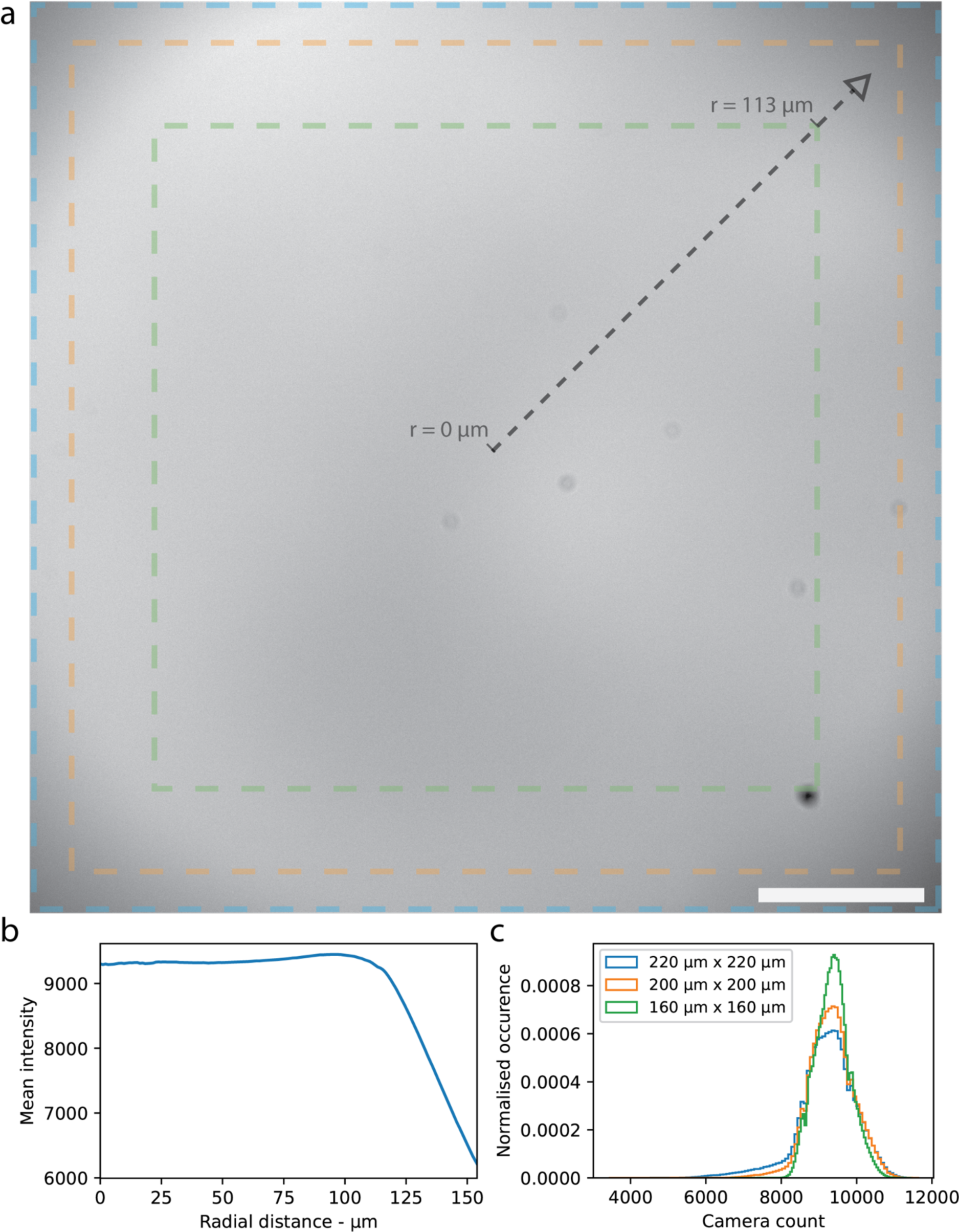
Vignetting limits the maximum homogeneous FOV. (a) ASTER excitation of a thin layer of Nile-Blue exhibiting fading at the edges. Scalebar, 40 µm. (b) Mean intensity at increasing distance from the center. (c) Intensity histogram for the full 220 µm x 220 µm field of the camera, a restricted 200 µm x 200 µm and 160 µm x 160 µm fields.

**Supplementary Figure 4:**
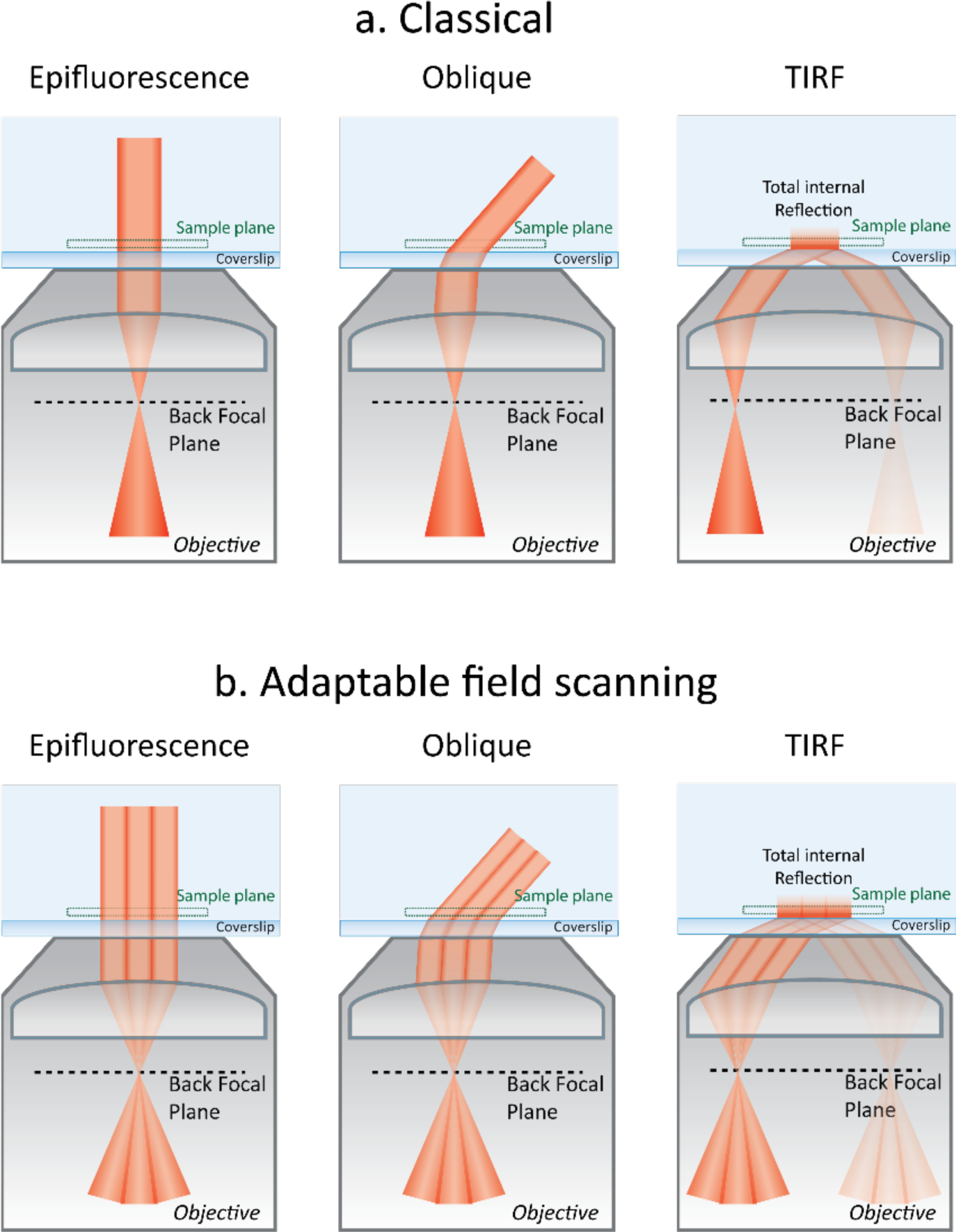
Implementation of TIRF and oblique illumination in classical and ASTER excitation schemes. (a) Classical configuration in EPI, oblique HiLo and TIRF, from left to right. Each position in the Back Focal Plane coincides with a given output angle. (b) ASTER configuration for EPI, oblique HiLo and TIRF, where the scanning effect modifies the effective field of view but does not affect the output angle.

**Supplementary Figure 5:**
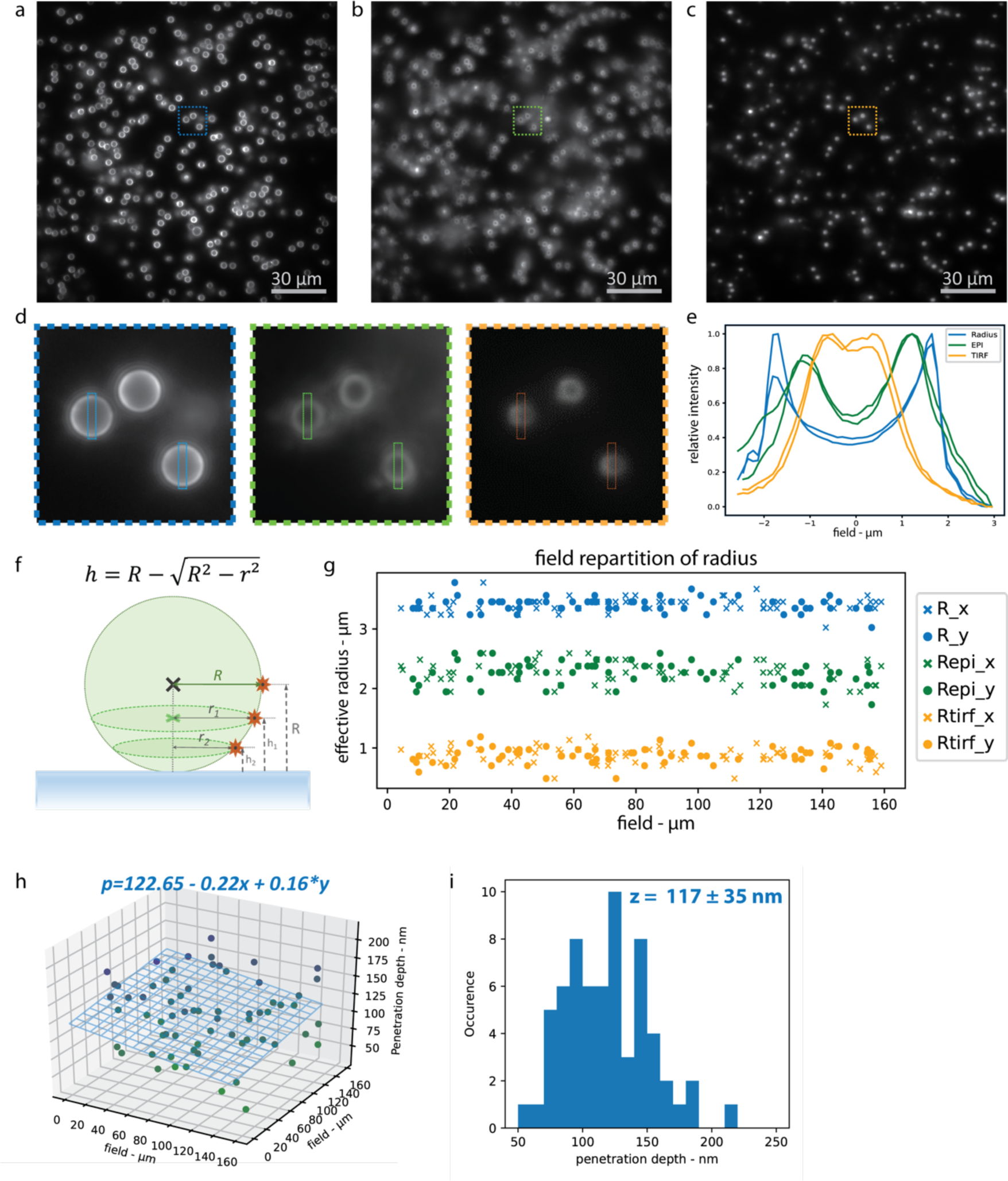
Measurement of the effective optical sectioning (excitation and detection) in epi and TIRF using labelled beads. (a) Imaging of beads in epi, with focus at the beads median planes. (b) Imaging of beads in epi, with focus at the coverslip. (c) Imaging of beads in TIRF, with focus at the coverslip. (d) Close up view of highlighted areas in (a-c) showing that each illumination condition results in its own effective bead radius. (e) Vertical profiles of highlighted cross-sections in (d). (f) Schematic of a nanobead, with R the median radius, and intermediary radii ri corresponding to different heights. (g) Distribution of the radius measured for each sphere along the field. A cross (respectively a circle) denotes a measurement along the x (respectively y) axis. (h) Fitting of the 2D distribution of penetration depth by a plane. (i) Distribution of the TIRF measured penetration depth.

**Supplementary Figure 6:**
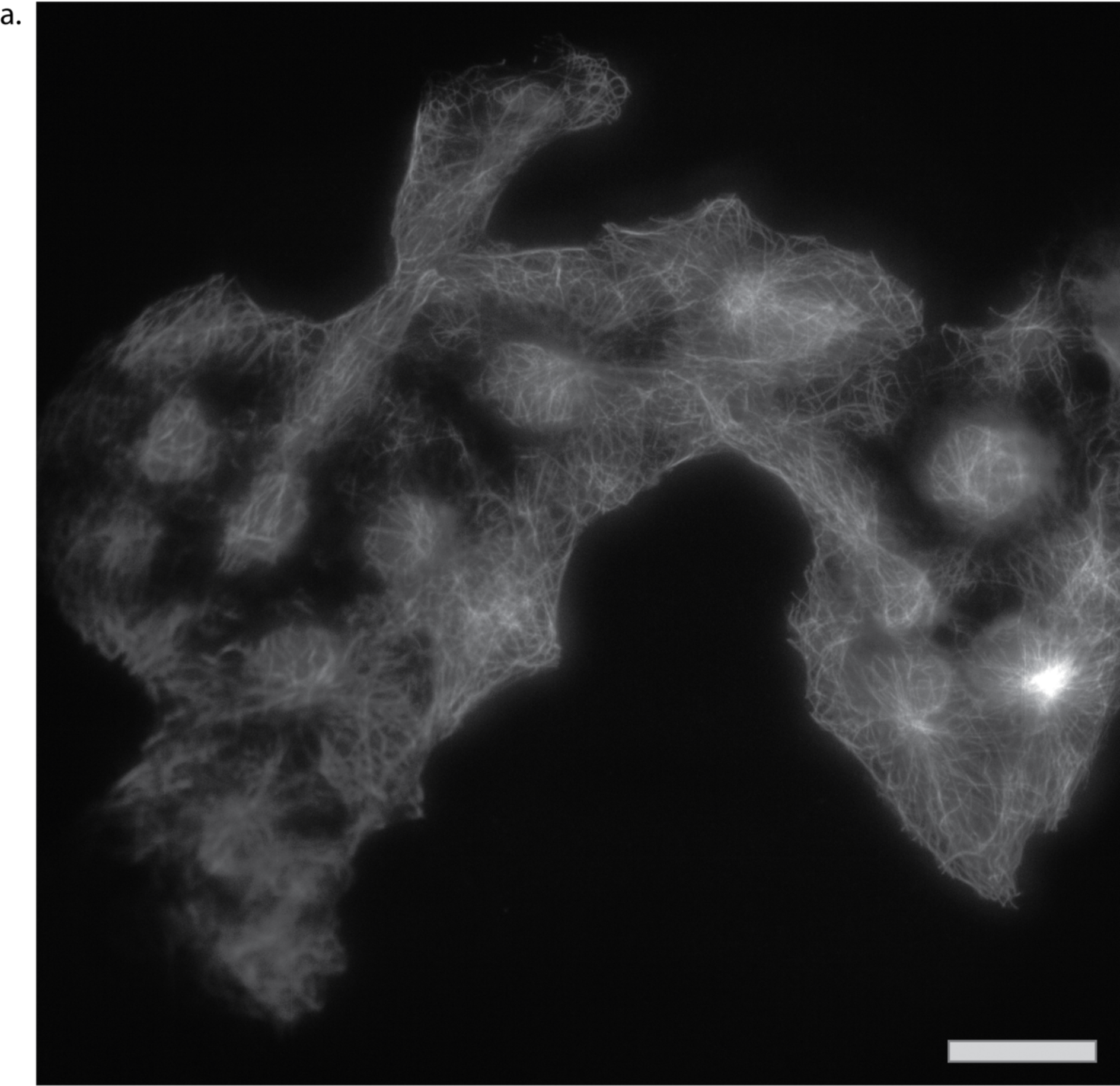
Full-field TIRF uniform ASTER excitation at 5 ms integration time. (a) Resulting image of COS-7 cells labeled for microtubules using AF647-coupled antibodies imaged over 220 µm x 220 µm. Scanning consisted in twelve lines scanned with a scanning period of 5 ms. Scalebar, 30 µm.

**Supplementary Figure 7:**
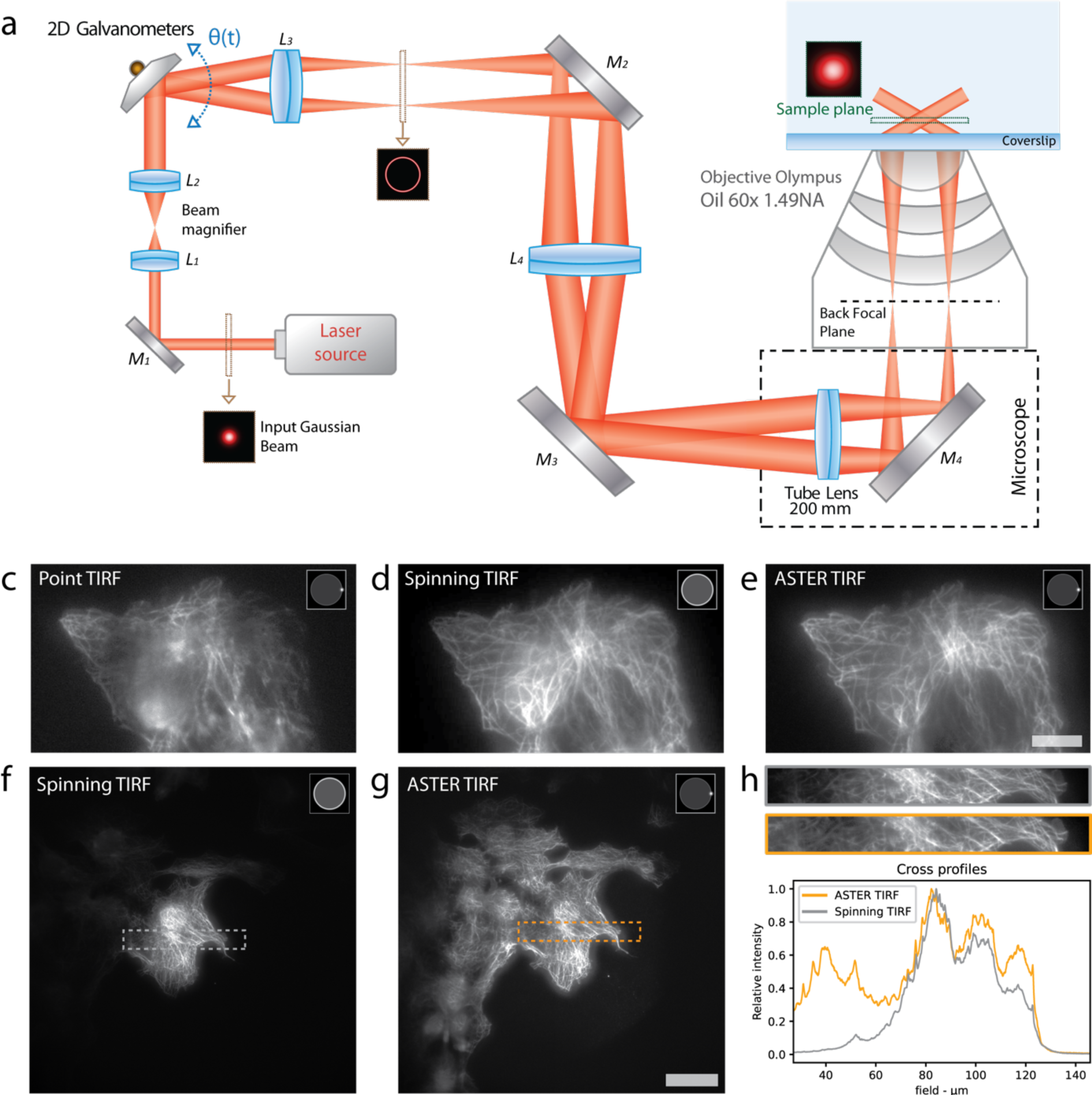
Homogeneity of ASTER TIRF excitation compared to azimuthal spinning TIRF. **(a)** Adaptation of ASTER setup to perform azimuthal spinning TIRF by scanning the beam in a sample-conjugated plane. **(c-g)** TIRF images of COS-7 cells labeled for microtubules using AF647-coupled antibodies excited either with an azimuthal spinning setup without scanning (c), with scanning (d,f), or with an ASTER uniform excitation setup (e,g). Notable inhomogeneities in image (c) are not present in images (d) and (e), which exhibit similar TIRF quality. Scalebar, 10 µm. (f) and (g) are full field images, whose highlighted sections are shown in (h). Compared to ASTER, the gray profile from image (f) show a regular decrease in intensity around 80 µm. Scalebar, 40 µm

**Supplementary Figure 8:**
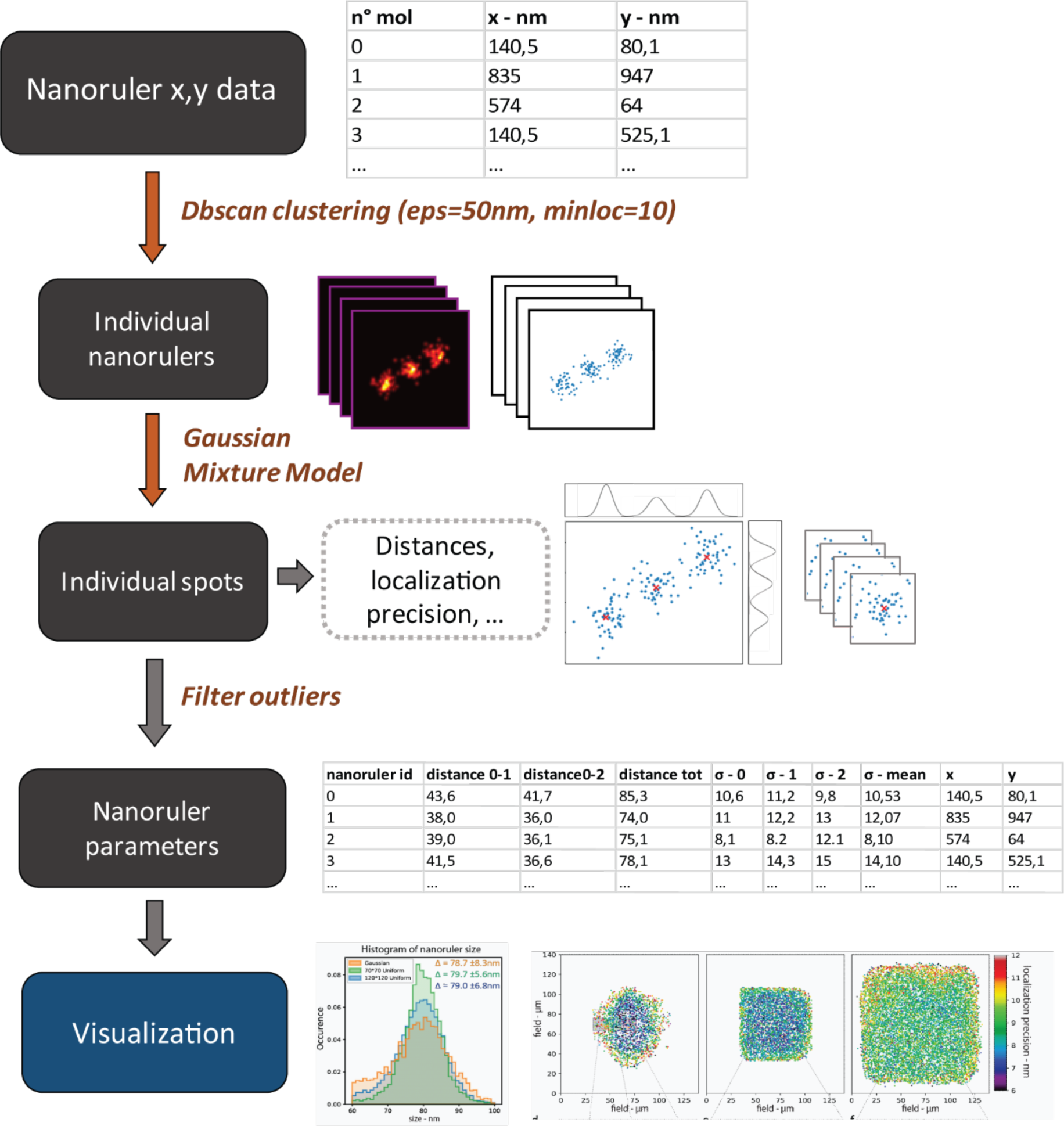
Workflow Analysis for three spot nanorulers. (See Methods). From the cloud of localization points, a DBscan isolates each individual nanoruler. One nanoruler consists of three aligned spots, each separated by 40 nm. For each individual nanoruler a gaussian mixture model fit the localization point clouds by three 2D normal distributions. This estimates which parameters are the most likely to produce the observed point distribution, namely the position and standard deviation of each of the three spots. Each point can then be associated to its most probable spot. This allows for measurement of nanoruler sizes, and estimation of localization precision for all individual nanorulers. Visualization is then performed with the Python library matplotlib.

**Supplementary Figure 9:**
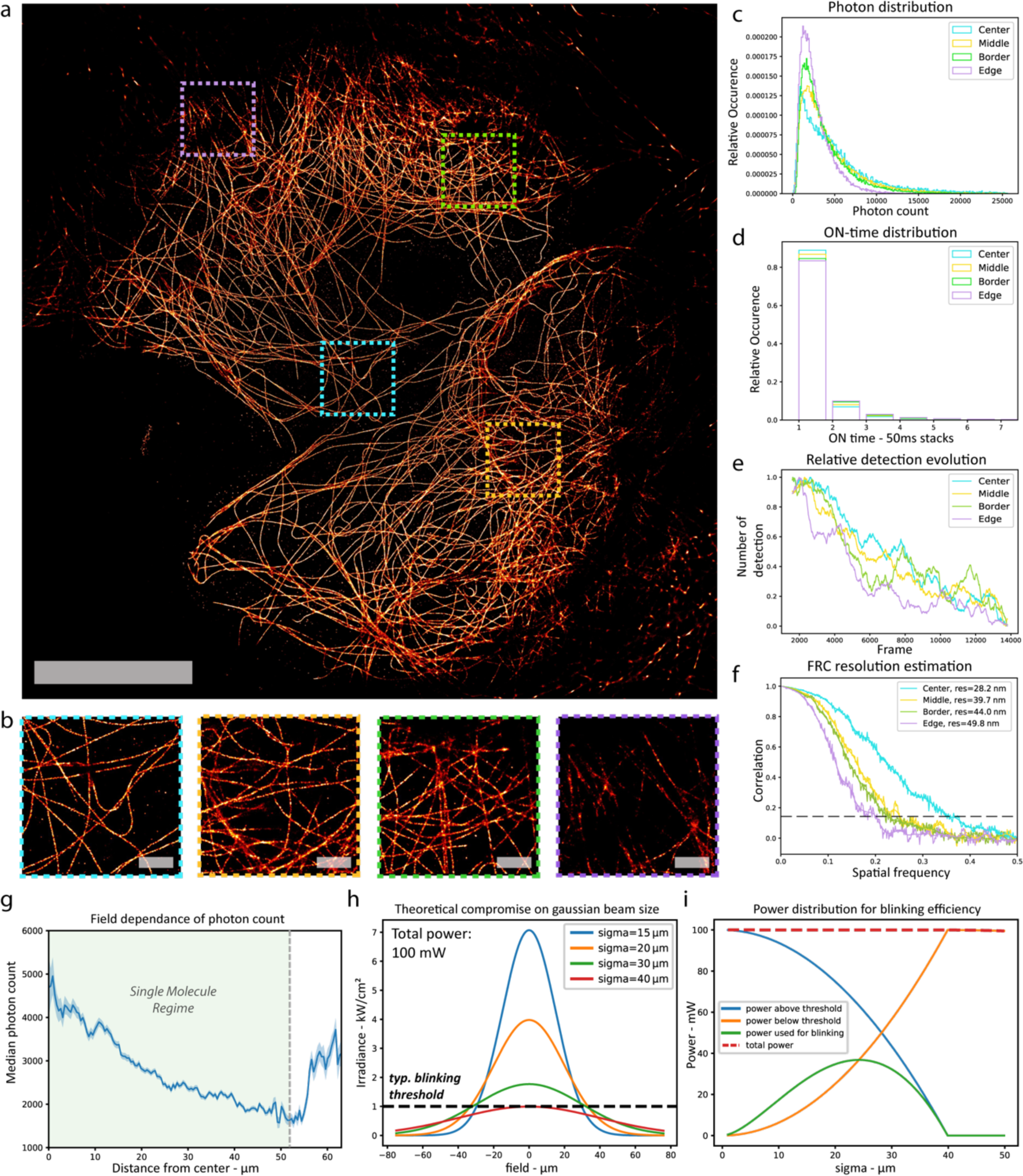
Gaussian illumination effects in single molecule STORM microscopy (σ = 45 µm). **(a-b)** Gaussian STORM imaging of COS-7 cells labeled for microtubules using AF647-coupled antibodies. Detection was done on 20000 frames at 50 ms exposure time with 300 mW laser power. Scalebar, 10 µm. (b) shows close up views of highlighted areas in (a). Scalebar 1µm. **(c)** Photon count distribution histogram for highlighted areas in (a). **(d)** Blinking ON-time distribution for highlighted areas in (a), expressed in number of frames (50 ms). **(e)** Temporal evolution of detection count for highlighted areas in (a). **(f)** FRC estimation of resolution for highlighted areas in (a). **(g)** Radial median photon count distribution. When the single molecule regime is broken, multiple fluorophores are detected and result in overestimation of photon count. **(h)** Compromise on gaussian beam size to reach a given blinking threshold with a fixed total power of 100 mW. **(i)** Resulting repartition of power usage. Power used for blinking is the product of the blinking threshold by the area that is illuminated over this threshold.

**Supplementary Figure 10:**
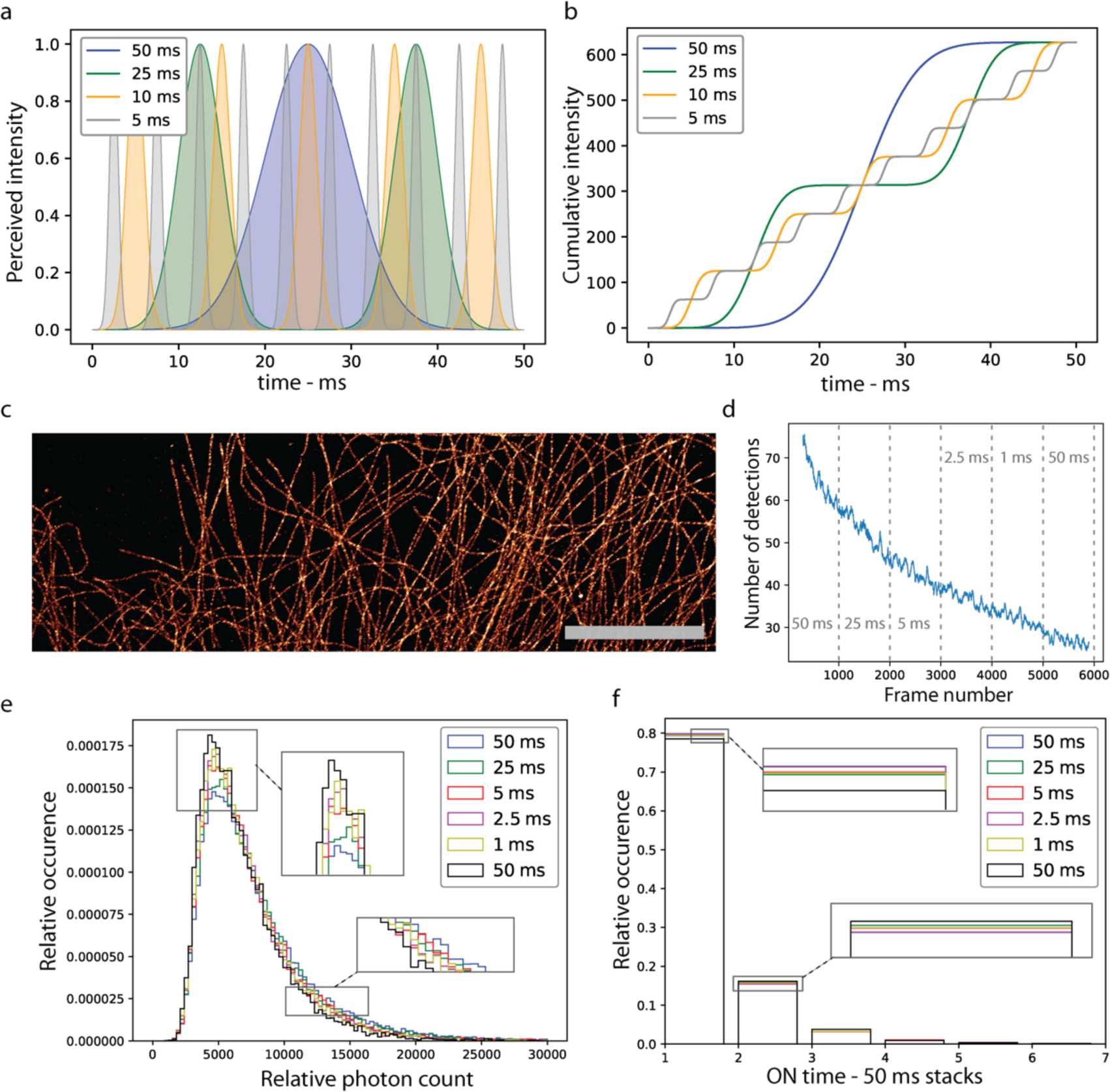
Impact of scanning on STORM blinking properties. **(a)** Perceived irradiance along time for a fluorophore excited under different ASTER scanning periods. **(b)** Cumulative intensity for each scanning period over 50 ms, resulting in a similar mean irradiance. **(c-f)** STORM imaging of COS-7 cells labeled for microtubules using AF647-coupled antibodies over a constant field with varying scanning periods. Scanning period is modified every 1000 frames and chronologically takes values 50, 25, 5, 2.5, 1 and 50 ms. (c) Resulting image over the whole frames. Scalebar, 10 µm. (d) Number of molecules detected per frame. Gray text indicates the experimental scanning speed for each frame range. (e) Relative photon count histogram for each scanning speed. (f) Relative ON time histogram for each scanning speed.

**Supplementary Figure 11:**
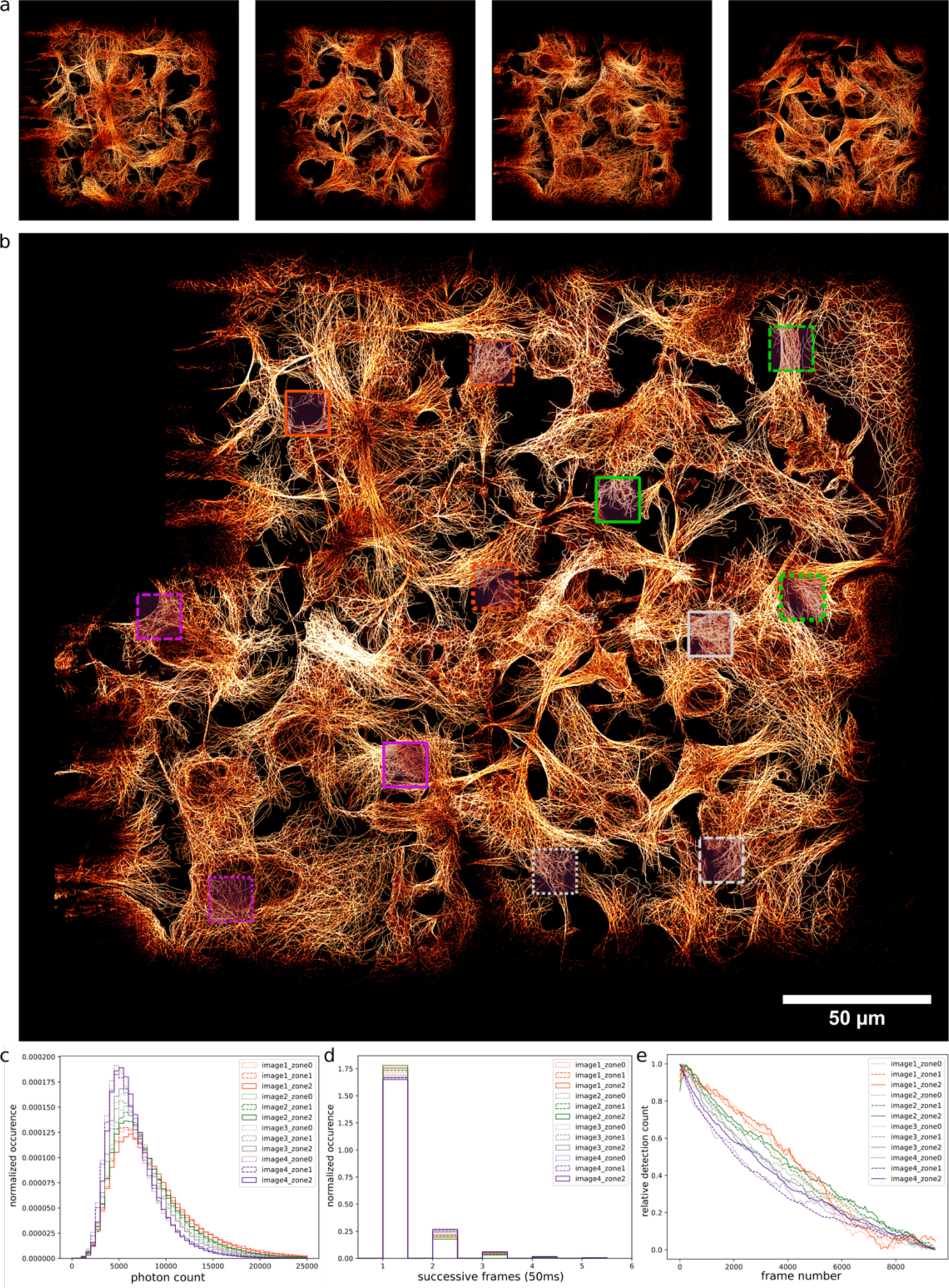
Stitching of four STORM images of COS-7 labeled for microtubules, resulting in a 300 µm x 300 μm hyper-large FOV. **(a)** Individual 150 µm x 150 μm STORM images. **(b)** Stitched images resulting in a 300 µm x 300 μm field of view. **(c-e)** Normalized photon count histograms (c), normalized ON time histograms (d) and relative number of detections along frames (e) for highlighted areas in (b).

**Supplementary Figure 12:**
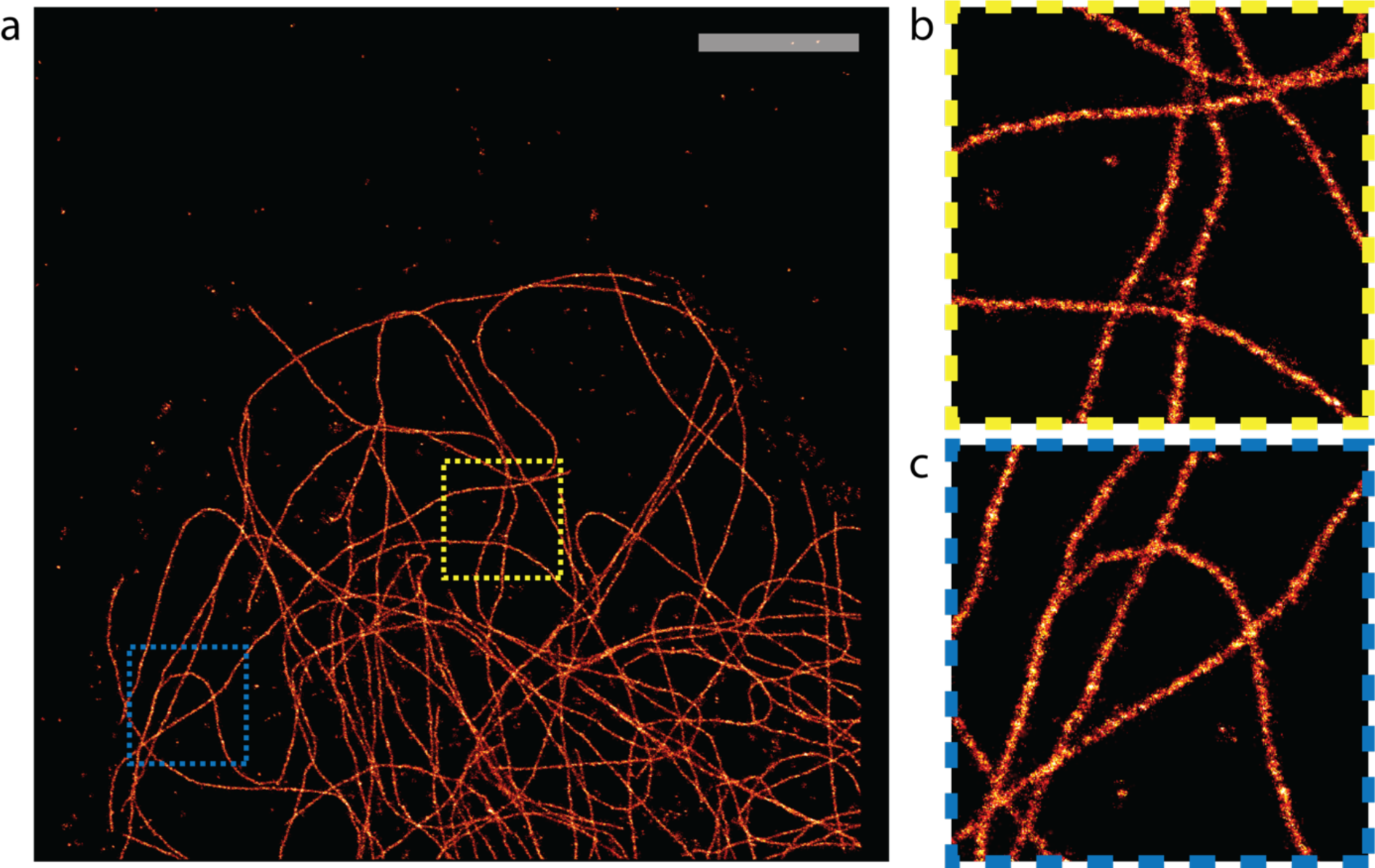
Fast STORM experiment acquired at 5ms integration time and 27 kW/cm& irradiance. **(a)** Resulting 10 nm-pixel image of COS-7 cells labeled for microtubules using AF647-coupled antibodies for a 20,000 frames (100 seconds) acquisition. After filtering outliers, 650,000 molecules contribute to the final image with an average density of 1000 molecules/µm^²^. **(b-c)** are close up views of highlighted areas in (a). Scalebar, 5 µm.

**Supplementary Figure 13:**
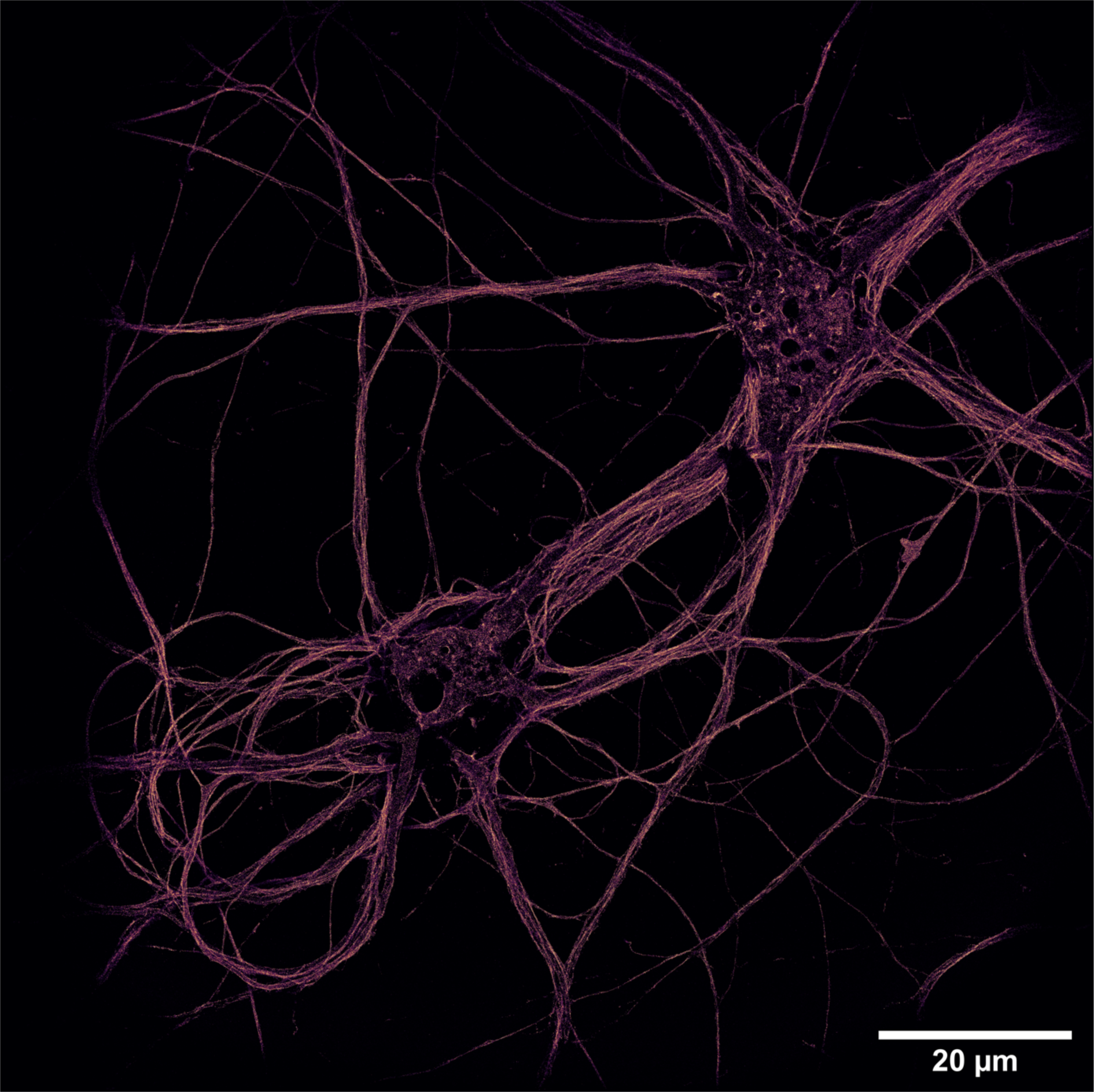
STORM 200 µm x 200 µm image of neuronal β2-spectrin, labeled with AF647.

**Supplementary Figure 14:**
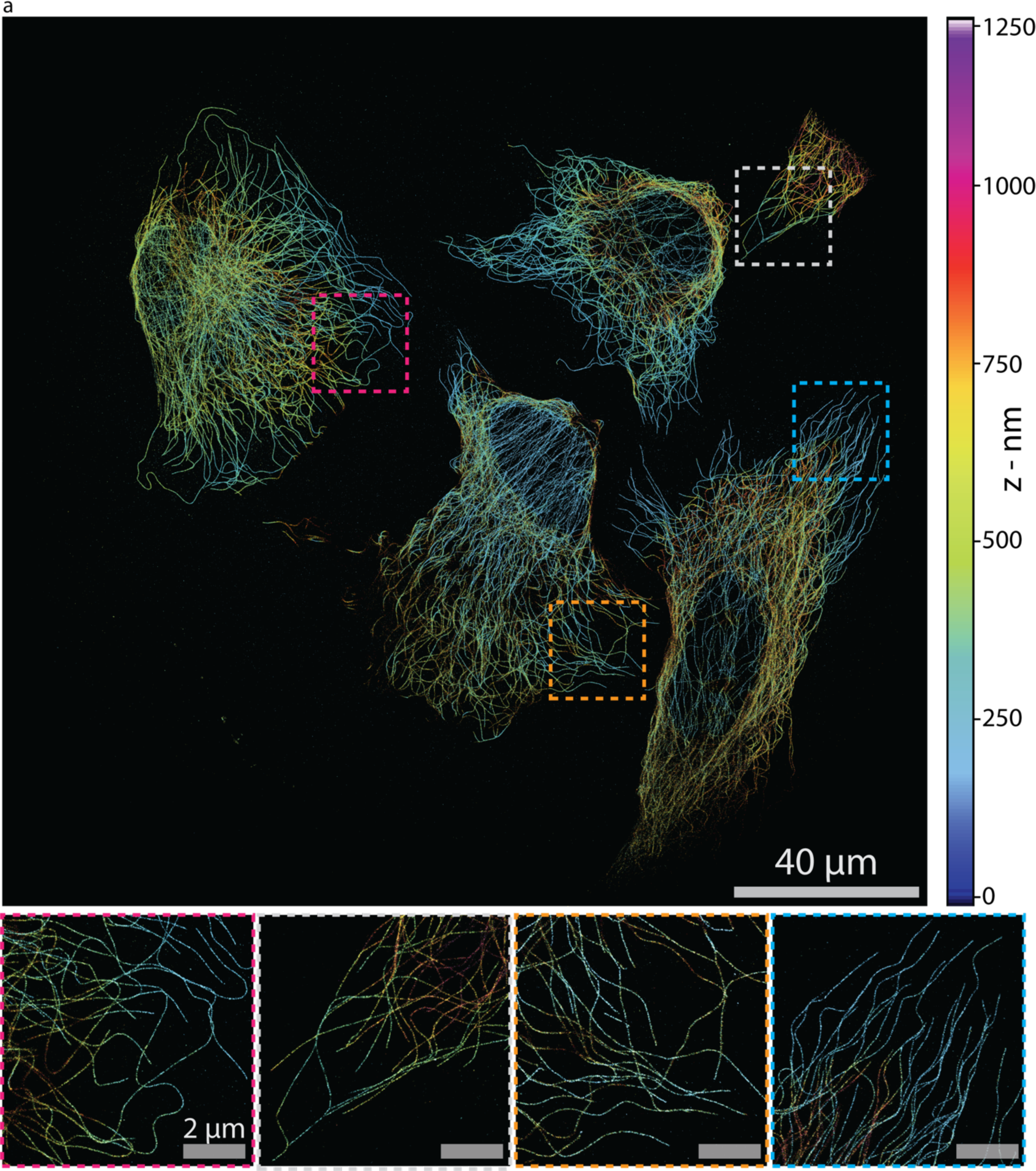
STORM 200 µm x 200 µm 3D image of COS-7 cells labeled for microtubules using AF647-coupled antibodies. **(a)** 3D color encoded image of microtubules, reconstruction pixel is 40nm and a 1-pixel gaussian blur has been applied. Axial information was obtained by splitting the detection relay in two paths and placing a 16k astigmatic lens on one path. **(b)** 20 µm x 20 µm images of highlighted areas in (a), with a reconstruction pixel of 10 nm.

**Supplementary Notes 1:** Comparison of uniform excitation methods

There are many ways to provide a uniform illumination, which can be classified in two general categories, and compared with ASTER:

### Waveguides

Waveguides provide uniformity on extremely large FOV, but are impossible to adapt as they provide a fixed excitation depth, and a fixed illumination size. They illuminate the sample with evanescent waves and are adapted to experiments that needs low power, with optical sectioning restrained to the proximity of the coverslip, as is the case for PAINT experiments. They are relatively low cost and achromatic, but may be complex to implement as mounting of the sample and coupling of the laser is not performed in a classical way.

### Direct beam reshapers

Beam reshapers directly transform a gaussian propagative beam into a flat-top. This includes optical beam-shapers (such as those from Pishaper, Topshape or Topag), square core fibers; microlens-array and phase SLM. Due to the nature of light, the flat-top profile is not maintained along its propagation and care must be given to the optical alignment of optical planes, so that the top-hat yields optimal contrast at sample plane. Beam reshapers are relatively achromatic, and with the addition of a translation stage allow shifting of the output illumination angle to perform optical sectioning techniques.

A primary drawback of direct beam-reshapers is that they are ill-adapted to quantitative TIRF: in the back focal plane, the flat-top profile should approximately take the shape of its Fourier transform, resulting in a sinc shape. Even though qualitative TIRF is achievable it should be hampered by multiple output angles, as it is not possible to precisely restrain the width of the beam in the back focal plane.

Furthermore, such system may suffer from speckle patterns. A rotating diffuser or vibrating membranes can be placed on the path of the beam and will typically average out the speckle in the order of 10 ms. When using fibers however, it is possible to use a vibrating motor to average the speckle in less than 1 ms, but will waste more laser power than beam-shapers elements as coupling in the fiber will typically waste 40% of the input power. It can be balanced by using multiple lasers with beam combiners, but will increase the system complexity and cost. In term of FOV size, beam reshapers can always be supplemented with an afocal system, so that even if the input and/or output beam is restrained in size, it will not be limiting in the global illumination setup, but once a size has been chosen it will be complex to change. While optical beam shapers and fibers will require physical intervention to adapt the flat-top size, the microlens-array device can slightly adapt the illumination by moving its components. On this matter, piSMLM, which uses a phase only SLM to shape the beam is extremely adaptable, but also results in great power losses (∼90%).

### ASTER

ASTER is an hybrid scanning-wide-field illumination setup and is close in performance to optical beam-reshapers. It continuously scans a gaussian beam following specific patterns to provide a flat-top in a time-averaged manner. With the galvanometer technology, one line can typically be scanned in 300µs, so when using the technology at its limit 16 lines can be scanned in 5 ms to generate a flat-top on wide fields. Furthermore, even though a flat-top is synthetized, the beam remains gaussian at each instant, and the synthetized field keeps its properties along prop-agation. In particular, TIRF should not be hampered given that the scanning device is well conjugated to the back focal plane of the objective, where the beam will take the shape of a focalized gaussian.

No speckle pattern was observed when illuminating with ASTER in EPI, HiLo or TIRF configurations. This is mostly attributed to the scanning effect, that means out the speckle as it simultaneously generates the flat-top and thus gives comparable results to azimuthal spinning TIRF. ASTER is not restrained in input beam size and can easily generate variable excitation sizes and positions. As it is controlled electronically, it can shift from one configuration to another in milliseconds. In that regard, ASTER is the most versatile of all methods, but comes with a complexity in time dependance. The scanning pattern should always be adapted so that the synthesis of uniform illumination does not yield unwanted stroboscopic effects. Assuring that the period of the flat-top synthesis is two times less than the period of the observed phenomena should allow for confident observation.

All of these properties are qualitatively synthetized in this table:

| General Scheme | Adaptability | Sectioning | Power Efficiency | Lowest image rate without speckle | Cost of devices | Ease of implementation |
| --- | --- | --- | --- | --- | --- | --- |
| **Waveguide** | poor | great, fixed | - | not limited | low | Sample handling, coupling in waveguide |
| **Multimode fiber** | physical intervention | average | average | fast motor - 0.5ms | 0.2 k€ * | Fiber coupling |
| **Optical Beam Shaper** | physical intervention | average | good | diffuser - several ms | 0.5-5 k€ | - |
| **Microlens array** | average | not shown | good | diffuser - several ms | 0.6k€ | Alignment |
| **phase SLM** | excellent | not shown | poor | diffuser - several ms | 10-15 k€ | Alignment, control of SLM, calibration |
| **ASTER** | excellent | good | good | 2ms - 5ms (FOV dependant) | 1-5 k€ | Alignment, electronic control |

\*actual cost for multimode fiber is higher when considering multiple lasers, beam couplers and rotating motor.

**Supplementary Notes 2:** Relation between minimum frame rate and field size with ASTER

ASTER relies on the continuous scan of a wide beam to generate a temporally-averaged flat-top profile. The minimum time needed to generate this profile depends on the size **σ** of the initial beam, and on the resulting side length **D** of the field.

As can be seen in Supplementary Figure 1, in order to generate a flat illumination the minimum distance between close lines must be less than 1.7σ. In this article we chose a minimal spacing of 1.2σ. Notably, the flat-top effect is maintained even when this length is diminished and scanning close lines should not impact fluorophore blinking (see Supplementary Fig. 10). A small gap however allows us to keep a flat-top profile even when reducing the size of the initial gaussian beam. Considering the fact than the step response of the galvanometers is around 300 µs, a line can be scanned in at least 300 µs. We then can estimate the minimum time necessary to generate a field of length **D**. For example, at least 11 lines are needed to generate a uniform field of size D=12σ, resulting in a 3.6 ms minimum generation time. The general formula is:

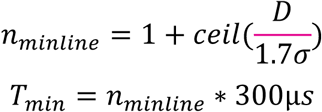

Notably, as we do not perform a point by point scanning but scans line, this relation is not proportional to the area of the field but to its longest length. This formula also shows that the wider the size of the initial beam, the faster we can generate a flat-top. However, the border of the flat-top will exhibit similar shape than that of the initial scanned beam, so that contrast will be hampered if the initial beam is chosen too wide.

In this publication, the initial beam size is 17 µm. Fields of 200 µm x 200 µm can then be generated by scanning 10 lines with a gap of 1.4 σ which can theoretically be achieved in 3 ms. Smaller fields, such as 30 µm x30 µm can be achieved with our beam size by scanning 3 lines in less than a millisecond.

In general for SMLM, integration times in the order of 20-100 ms are used, so there is no need to achieve the fastest scanning speed. Instead, the period of the field synthesis should be around two to four times less that of the integration time, this will be enough to guarantee a correct mean flat-top and maintain the galvanometers in a favorable regime.

**Supplementary Notes 3:** Uncertainties in measurement of microbead excitation depth Calibration of the TIRF penetration depth among microbeads yields a 115 nm mean value with a standard deviation of 35 nm (Supplementary Figure 3). This arises from the calculus of radii, which have a standard deviation of 0.11µm for the median radius, and 0.14 µm for the TIRF effective radius. This radius deviation corresponds to a precision of approximately one pixel (108 nm) and partly reflects the profiles that are represented in Supplementary Figure 3.e, where some peaks cannot confidently be attributed between two close pixels. It is plausible that smaller pixel size would improve precision. However, assuming that we regularly miss the position of peaks by 0.5 pixel, the deviation should be around 54 nm and not 108 nm. Other sources of variation should be taken into account, as the sample is not perfect: there may be flattening of some microbeads, inhomogeneities at the coverslip and/or a spatial tilt of the sample.

We fitted the measured penetration depth along the field by a plane, and found that a 1µm deviation in the x direction (respectively y) shifted the measured mean penetration depth by -0.22 nm (respectively 0.16 nm), When taking this tilt into account the deviation of the penetration depth would be of 33.0 nm, which indicates that this tilt does not greatly contributes to the deviation. This tilt could be indicative of either tilt of the coverslip, or slight misalignment of the galvanometers with the back focal plane of the objective. However, in the latter case, a dependence between field and penetration depth would arise in the direction of the incidence angle (namely x), but not in both directions.

Finally, we checked for local spatial correlation by calculating Moran I and Geary C indexes. Theses indexes characterize the spatial correlation between measurements by using weights as indicators of proximity. For Moran I index, positive spatial correlation (respectively negative) is indicated by a value close to 1 (respectively -1). For Geary C index, positive spatial correlation (respectively negative) is indicated by a value close to 0 (respectively 2).

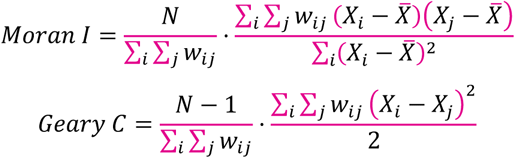

The choice of the weight factors **wij** will determine the final results. In order to conclude objectively we calculate Moran and Geary indexes using both boolean and continuous weights. Boolean weights **wij** equal 1 if and only if beads i and j are the closest neighbors, while continuous weights are based on distances and increase with proximity between beads.

| Indexes | Coefficients |  |
| --- | --- | --- |
|  | Continuous | Booleans |
| Moran I | -0.004 | 0.245 |
| Geary C | 0.996 | 0.843 |

Table of the Calculated Moran I and Geary C indexes for n=66 microbeads from Figure 3. Weights were ether chosen continuously with distances or as Booleans with closest neighbour.

As can be assessed the above table, values of indexes are close to 0 for Moran, and 1 for Geary C indexes, no matter the choice of weights. This is indicative of a random spatial distribution at the scale of the microbeads. We conclude that there is no strong spatial correlation between close beads (micrometers apart) and their measured penetration depth. Though local inhomogeneities may exist at smaller scales, the effective optical sectioning of our experiment can be considered globally uniform.

## References

1. Tokunaga, M., Imamoto, N. & Sakata-Sogawa, K. Highly inclined thin illumination enables clear singlemolecule imaging in cells. Nat. Methods 5, 159–161 (2008).

2. Nakata, T. & Hirokawa, N. Microtubules provide directional cues for polarized axonal transport through interaction with kinesin motor head. J. Cell Biol. 162, 1045–1055 (2003).

3. Guo, Y. et al. Visualizing Intracellular Organelle and Cytoskeletal Interactions at Nanoscale Resolution on Millisecond Timescales. Cell 175, 1430-1442.e17 (2018).

4. Stout, A. L. & Axelrod, D. Evanescent field excitation of fluorescence by epi-illumination microscopy. Appl. Opt. 28, 5237 (1989).

5. Axelrod, D. Cell-substrate contacts illuminated by total internal reflection fluorescence. J. Cell Biol. 89, 141–145 (1981).

6. Stock, K. et al. Variable-angle total internal reflection fluorescence microscopy (VA-TIRFM): realization and application of a compact illumination device. J. Microsc. 211, 19–29 (2003).

7. Konopka, C. A. & Bednarek, S. Y. Variable-angle epifluorescence microscopy: a new way to look at protein dynamics in the plant cell cortex. Plant J. 53, 186–196 (2008).

8. van ‘t Hoff, M., de Sars, V. & Oheim, M. A programmable light engine for quantitative single molecule TIRF and HILO imaging. Opt. Express 16, 18495 (2008).

9. Mattheyses, A. L., Shaw, K. & Axelrod, D. Effective elimination of laser interference fringing in fluorescence microscopy by spinning azimuthal incidence angle. Microsc. Res. Tech. 69, 642–647 (2006).

10. Fiolka, R., Belyaev, Y., Ewers, H. & Stemmer, A. Even illumination in total internal reflection fluorescence microscopy using laser light. Microsc. Res. Tech. 71, 45–50 (2008).

11. Boulanger, J. et al. Fast high-resolution 3D total internal reflection fluorescence microscopy by incidence angle scanning and azimuthal averaging. Proc. Natl. Acad. Sci. 111, 17164–17169 (2014).

12. Schreiber, B., Elsayad, K. & Heinze, K. G. Axicon based Bessel beams for flat-field illumination in total internal reflection fluorescence microscopy. Opt. Lett. 42, 3880 (2017).

13. Almada, P., Culley, S. & Henriques, R. PALM and STORM: Into large fields and high-throughput microscopy with sCMOS detectors. Methods 88, 109–121 (2015).

14. Betzig, E. et al. Imaging Intracellular Fluorescent Proteins at Nanometer Resolution. Science 313, 1642–1645 (2006).

15. Hess, S. T., Girirajan, T. P. K. & Mason, M. D. Ultra-High Resolution Imaging by Fluorescence Photoactivation Localization Microscopy. Biophys. J. 91, 4258–4272 (2006).

16. Sharonov, A. & Hochstrasser, R. M. Wide-field sub-diffraction imaging by accumulated binding of diffusing probes. Proc. Natl. Acad. Sci. 103, 18911–18916 (2006).

17. Rust, M. J., Bates, M. & Zhuang, X. Sub-diffraction-limit imaging by stochastic optical reconstruction microscopy (STORM). Nat. Methods 3, 793–796 (2006).

18. Heilemann, M. et al. Subdiffraction-Resolution Fluorescence Imaging with Conventional Fluorescent Probes. Angew. Chem. Int. Ed. 47, 6172–6176 (2008).

19. Heilemann, M., van de Linde, S., Mukherjee, A. & Sauer, M. Super-Resolution Imaging with Small Organic Fluorophores. Angew. Chem. Int. Ed. 48, 6903–6908 (2009).

20. Ramachandran, S., Cohen, D. A., Quist, A. P. & Lal, R. High performance, LED powered, waveguide based total internal reflection microscopy. Sci. Rep. 3, 2133 (2013).

21. Diekmann, R. et al. Chip-based wide field-of-view nanoscopy. Nat. Photonics 11, 322–328 (2017).

22. Archetti, A. et al. Waveguide-PAINT offers an open platform for large field-of-view super-resolution imaging. Nat. Commun. 10, 1267 (2019).

23. Douglass, K. M., Sieben, C., Archetti, A., Lambert, A. & Manley, S. Super-resolution imaging of multiple cells by optimized flat-field epi-illumination. Nat. Photonics 10, 705–708 (2016).

24. Khaw, I. et al. Flat-field illumination for quantitative fluorescence imaging. Opt Express 26, 15276–15288 (2018).

25. Rowlands, C. J., Ströhl, F., Ramirez, P. P. V., Scherer, K. M. & Kaminski, C. F. Flat-Field Super-Resolution Localization Microscopy with a Low-Cost Refractive Beam-Shaping Element. Sci. Rep. 8, 5630 (2018).

26. Stehr, F., Stein, J., Schueder, F., Schwille, P. & Jungmann, R. Flat-top TIRF illumination boosts DNA-PAINT imaging and quantification. Nat. Commun. 10, 1268 (2019).

27. Zhao, Z., Xin, B., Li, L. & Huang, Z.-L. High-power homogeneous illumination for super-resolution localization microscopy with large field-of-view. Opt. Express 25, 13382–13395 (2017).

28. Deschamps, J., Rowald, A. & Ries, J. Efficient homogeneous illumination and optical sectioning for quantitative single-molecule localization microscopy. Opt. Express 24, 28080 (2016).

29. Kwakwa, K. et al. easySTORM: a robust, lower-cost approach to localisation and TIRF microscopy. J. Biophotonics 9, 948–957 (2016).

30. Chen, S.-Y., Bestvater, F., Schaufler, W., Heintzmann, R. & Cremer, C. Patterned illumination single molecule localization microscopy (piSMLM): user defined blinking regions of interest. Opt. Express 26, 30009 (2018).

31. Kurvits, J. A., Jiang, M. & Zia, R. Comparative analysis of imaging configurations and objectives for Fourier microscopy. J. Opt. Soc. Am. A 32, 2082 (2015).

32. Mattheyses, A. L. & Axelrod, D. Direct measurement of the evanescent field profile produced by objectivebased total internal reflection fluorescence. J. Biomed. Opt. 11, 014006 (2006).

33. Cabriel, C., Bourg, N., Dupuis, G. & Lévêque-Fort, S. Aberration-accounting calibration for 3D single-molecule localization microscopy. Opt. Lett. 43, 174 (2018).

34. Ester, M., Kriegel, H.-P., Sander, J. & Xu, X. A density-based algorithm for discovering clusters in large spatial databases with noise. in 226–231 (AAAI Press, 1996).

35. Tam, J., Cordier, G. A., Borbely, J. S., Sandoval Ál-varez, Á. & Lakadamyali, M. Cross-Talk-Free MultiColor STORM Imaging Using a Single Fluorophore. PLoS ONE 9, e101772 (2014).

36. Xu, J., Ma, H. & Liu, Y. Stochastic Optical Reconstruction Microscopy (STORM). Curr. Protoc. Cytom. 81, (2017).

37. Lin, Y. et al. Quantifying and Optimizing Single-Mol-ecule Switching Nanoscopy at High Speeds. PLOS ONE 10, e0128135 (2015).

38. Banterle, N., Bui, K. H., Lemke, E. A. & Beck, M. Fourier ring correlation as a resolution criterion for superresolution microscopy. J. Struct. Biol. 183, 363–367 (2013).

39. Bates, M., Blosser, T. R. & Zhuang, X. Short-Range Spectroscopic Ruler Based on a Single-Molecule Optical Switch. Phys. Rev. Lett. 94, 108101 (2005).

40. Diekmann, R. et al. Optimizing imaging speed and excitation intensity for single-molecule localization microscopy. Nat. Methods (2020) doi:10.1038/s41592-020-0918-5.

41. Heuser, J. Three-dimensional visualization of coated vesicle formation in fibroblasts. J. Cell Biol. 84, 560–583 (1980).

42. Lampe, M., Vassilopoulos, S. & Merrifield, C. Clathrin coated pits, plaques and adhesion. J. Struct. Biol. 196, 48–56 (2016).

43. Leterrier, C. et al. Nanoscale Architecture of the Axon Initial Segment Reveals an Organized and Robust Scaffold. Cell Rep. 13, 2781–2793 (2015).

44. Ganguly, A. et al. A dynamic formin-dependent deep F-actin network in axons. J. Cell Biol. 210, 401–417 (2015).

45. Xu, K., Zhong, G. & Zhuang, X. Actin, Spectrin, and Associated Proteins Form a Periodic Cytoskeletal Structure in Axons. Science 339, 452–456 (2013).

46. Leterrier, C., Dubey, P. & Roy, S. The nano-architecture of the axonal cytoskeleton. Nat. Rev. Neurosci. 18, 713–726 (2017).

47. Vassilopoulos, S., Gibaud, S., Jimenez, A., Caillol, G. & Leterrier, C. Ultrastructure of the axonal periodic scaffold reveals a braid-like organization of actin rings. Nat. Commun. 10, 5803 (2019).

48. Potsaid, B., Bellouard, Y. & Wen, J. T. (ASOM): A multidisciplinary optical microscope design for large field of view and high. 15 (2005).

49. Potsaid, B., Finger, F. P. & Wen, J. T.-Y. Living organism imaging with the adaptive scanning optical microscope (ASOM). in (eds. Farkas, D. L., Leif, R. C. & Nicolau, D. V.) 64411D (2007). doi:10.1117/12.699552.

50. Huang, B., Wang, W., Bates, M. & Zhuang, X. Three-Dimensional Super-Resolution Imaging by Stochastic Optical Reconstruction Microscopy. Science 319, 810–813 (2008).

51. Cabriel, C. et al. Combining 3D single molecule localization strategies for reproducible bioimaging. Nat. Commun. 10, 1980 (2019).

52. Cnossen, J. et al. Localization microscopy at doubled precision with patterned illumination. Nat. Methods 17, 59–63 (2020).

53. Reymond, L. et al. SIMPLE: Structured illumination based point localization estimator with enhanced preci-sion. Opt. Express 27, 24578 (2019).

54. Eilers, Y., Ta, H., Gwosch, K. C., Balzarotti, F. & Hell, S. W. MINFLUX monitors rapid molecular jumps with superior spatiotemporal resolution. Proc. Natl. Acad. Sci. 115, 6117–6122 (2018).

55. Jouchet, P. et al. Nanometric axial localization of single fluorescent molecules with modulated excitation. http://biorxiv.org/lookup/doi/10.1101/865865 (2019) doi:10.1101/865865.

56. Bossi, M. et al. Multicolor Far-Field Fluorescence Nanoscopy through Isolated Detection of Distinct Molecular Species. Nano Lett. 8, 2463–2468 (2008).

57. Fricke, F., Beaudouin, J., Eils, R. & Heilemann, M. One, two or three? Probing the stoichiometry of membrane proteins by single-molecule localization microscopy. Sci. Rep. 5, 14072 (2015).

58. Hung, Y.-J., Chang, H.-J., Chang, P.-C., Lin, J.-J. & Kao, T.-C. Employing refractive beam shaping in a Lloyd’s interference lithography system for uniform periodic nanostructure formation. J. Vac. Sci. Technol. B Nanotechnol. Microelectron. Mater. Process. Meas. Phenom. 35, 030601 (2017).

59. Weber, D. et al. Use of beam-shaping optics for wafer-scaled nanopatterning in laser interference lithography. Appl. Phys. A 125, 307 (2019).

60. Ishikawa-Ankerhold, H., Ankerhold, R. & Drummen, G. Fluorescence Recovery After Photobleaching (FRAP). in eLS (ed. John Wiley & Sons Ltd) a0003114 (John Wiley & Sons, Ltd, 2014). doi:10.1002/9780470015902.a0003114.

61. Dreier, J. et al. Smart scanning for low-illumination and fast RESOLFT nanoscopy in vivo. Nat. Commun. 10, 556 (2019).

62. Kaech, S. & Banker, G. Culturing hippocampal neurons. Nat. Protoc. 1, 2406–2415 (2006).

63. Jimenez, A., Friedl, K. & Leterrier, C. About samples, giving examples: Optimized Single Molecule Localization Microscopy. Methods 174, 100–114 (2020).

64. Bourg, N. et al. Direct optical nanoscopy with axially localized detection. Nat. Photonics 9, 587–593 (2015).

65. Feret, L. La grosseur des grains des matières pulvérulentes. (1930).

